# Intracellular *Staphylococcus aureus* employs the cysteine protease staphopain A to induce host cell death in epithelial cells

**DOI:** 10.1101/2020.02.10.936575

**Authors:** Kathrin Stelzner, Tobias Hertlein, Aneta Sroka, Adriana Moldovan, Kerstin Paprotka, David Kessie, Helene Mehling, Jan Potempa, Knut Ohlsen, Martin J. Fraunholz, Thomas Rudel

## Abstract

*Staphylococcus aureus* is a major human pathogen, which can invade and survive in non-professional and professional phagocytes. Intracellularity is thought to contribute to pathogenicity and persistence of the bacterium. Upon internalization by epithelial cells, cytotoxic *S. aureus* strains can escape from the phagosome, replicate in the cytosol and induce host cell death. Here, we identified a staphylococcal cysteine protease to induce cell death by intracellular *S. aureus* after translocation into the host cell cytoplasm. We demonstrated that loss of staphopain A function leads to delayed onset of host cell death and prolonged intracellular replication of *S. aureus* in epithelial cells. Overexpression of staphopain A in a non-cytotoxic strain facilitated intracellular killing of the host cell even in the absence of detectable intracellular replication. Moreover, staphopain A contributed to efficient colonization of the lung in a mouse pneumonia model. Our study suggests that staphopain A is utilized by *S. aureus* to mediate escape from the host cell and thus contributes to tissue destruction and dissemination of infection.

**Author Summary:** *Staphylococcus aureus* is a well-known antibiotic-resistant pathogen that emerges in hospital and community settings and can cause a variety of diseases ranging from skin abscesses to lung inflammation and blood poisoning. The bacterium asymptomatically colonizes the upper respiratory tract and skin of about one third of the human population and takes advantage of opportune conditions, like immunodeficiency or breached barriers, to cause infection. Although *S. aureus* is not regarded as a professional intracellular bacterium, it can be internalized by human cells and subsequently exit the host cells by induction of cell death, which is considered to cause tissue destruction and spread of infection. The bacterial virulence factors and underlying molecular mechanisms involved in the intracellular lifestyle of *S. aureus* remain largely unknown. We identified a bacterial cysteine protease to contribute to host cell death mediated by intracellular *S. aureus*. Staphopain A induced killing of the host cell after translocation of the pathogen into the cell cytosol, while bacterial proliferation was not required. Further, the protease enhanced survival of the pathogen during lung infection. These findings reveal a novel, intracellular role for the bacterial protease staphopain A.

## Introduction

*Staphylococcus aureus* is a Gram-positive bacterium frequently colonizing human skin and soft tissue, primarily the anterior nares, as part of the normal microflora [1]. However, in hospital- as well as in community-settings it arises as an opportunistic pathogen causing a plethora of diseases ranging from local, superficial skin infections, wound infections and abscesses to invasive, systemic diseases like osteomyelitis, pneumonia, endocarditis or sepsis [2]. This can be largely attributed to its vast array of virulence factors [3]. In addition, the emergence and rapid spread of methicillin-resistant *S. aureus* (MRSA) strains makes this pathogen particularly difficult to treat and leads to significant morbidity and mortality worldwide [4].

Whereas *S. aureus* is originally considered an extracellular pathogen, substantial evidence exists that it is able to invade non-phagocytic mammalian cells, like epithelial and endothelial cells, osteoblasts, fibroblasts or keratinocytes [e.g. 5, 6-8], as well as to survive internalization by professional phagocytes [e.g. 9, 10, 11]. Several studies demonstrate the existence of intracellular *S. aureus* in tissue and phagocytic cells *in vivo* [e.g. 12, 13-16]. Invasion of tissue cells is facilitated by numerous different bacterial adhesins and followed by escape from the bacteria-containing vacuole and cytosolic replication [reviewed in 17, 18, 19]. In professional phagocytes, intracellular *S. aureus* is able to resist the antimicrobial attack by the host cell and replication occurs within phagosomes. In both cell types, the pathogen eventually kills the host cell from within and a new infection cycle can be initiated. Numerous rounds of internalization and release may lead to excessive cell and tissue destruction, persistence and dissemination of infection, immune evasion and protection from antibiotic treatment [20–22].

Intracellular pathogens may induce killing as a strategy to exit from the host cell. Induction of programmed cell death, such as apoptosis, necroptosis or pyroptosis, or the damaging of host cell-derived membranes such as endosomal, vacuolar and plasma membrane have been demonstrated as mechanisms [reviewed in 23]. Reports on the molecular mechanisms underlying *S. aureus* induced host cell killing are rather inconsistent, which likely arises from the diversity of virulence factors within *S. aureus*. Some studies describe induction of apoptosis in professional as well as non-professional phagocytes [e.g. 24, 25-27], while others observed hallmarks of both apoptosis and necrosis [11, 28]. In epithelial cells, an autophagy-associated cell death was discovered [29], whereas in primary human polymorphonuclear neutrophils (PMNs) intracellular *S. aureus* induced a necroptotic cell death [30, 31]. The existence of different types of host cells and a multitude of different *S. aureus* strains further impedes the elucidation of the process and bacterial virulence factors involved therein.

To date, several *S. aureus* virulence factors have been linked to intracellular cytotoxicity. In non-professional phagocytes the hemolytic α-toxin was identified as a key factor mediating intracellular cytotoxicity [32, 33], whereas the bi-component leukotoxin LukAB (also known as LukGH) was shown to induce cell lysis after uptake of *S. aureus* by professional phagocytes [34–37]. Some reports also connected Panton-Valentine-leukotoxin (PVL) [35, 38, 39] or phenol-soluble modulins (PSMs) [40, 41] with host cell killing by intracellular *S. aureus*.

Beside these toxins, *S. aureus* secretes several proteases which were shown to contribute to virulence of the pathogen [42] and whose role in the intracellular lifestyle of the pathogen has not been investigated so far. *S. aureus* possesses two papain-like cysteine proteases, staphopain A (ScpA) and staphopain B (SspB), which have almost identical three-dimensional structures, despite sharing limited primary sequence identity [43, 44]. Both proteases are highly conserved among *S. aureus* isolates [45] and are expressed in the respective operons, *scp*AB and *ssp*ABC, together with their endogenous inhibitors, the staphostatins ScpB and SspC, which protect the bacteria from proteolytic degradation [46–48]. Staphopain A is secreted as zymogen and activated by autolytic cleavage once outside the bacterial cell [49]. By contrast, activation of staphopain B is the result of a proteolytic cascade initiated by aureolysin-mediated cleavage and activation of the V8 protease (SspA) which in turn processes SspB [47, 50]. *In vitro* experiments revealed a very broad activity of both enzymes on the destruction of connective tissue, the evasion of host immunity and the modulation of biofilm integrity [51–57]. A staphopain A-like protease with similar functions, called EcpA, is also expressed by *S. epidermidis* and other coagulase-negative staphylococci [42, 58, 59]. In contrast, orthologues of the *ssp*ABC operon have only been identified in *S. warneri* [60, 61].

In this study, we identified a novel role of *S. aureus* extracellular cysteine proteases in the intracellular lifestyle of the pathogen. We demonstrate that staphopain A, but not staphopain B, plays a role in intracellular killing of the host cell after phagosomal escape of *S. aureus*. Loss of function of staphopain A in a cytotoxic *S. aureus* strain led to delayed onset of host cell death in epithelial cells. Inducible expression of the cysteine protease in a non-cytotoxic bacterial strain initiated an apoptotic mode of cell death in the infected host cells, which was dependent on the protease activity. Additionally, staphopain A favored *S. aureus* colonization of the murine lung.

## Results

### Loss of function of a cysteine protease renders intracellular *S. aureus* less cytotoxic

In order to investigate whether *S. aureus* extracellular cysteine proteases play a role in induction of host cell death by intracellular bacteria, HeLa cells infected with the community-acquired MRSA (CA-MRSA) strain JE2 were treated with E-64d, a cell-permeable irreversible inhibitor of cysteine proteases. One hour post-infection all extracellular bacteria were removed by lysostaphin treatment and cell death was determined six hours post-infection by measurement of lactate dehydrogenase (LDH) release [62]. Addition of E-64d prior to infection led to a significant reduction (*p*=0.0022) in LDH release compared to cells treated with solvent control (DMSO) (Fig 1A). This observation led to the hypothesis that either the two extracellular cysteine proteases of *S. aureus*, staphopain A (*scp*A) or staphopain B (*ssp*B), or host cell cysteine proteases might be involved in *S. aureus* cytotoxicity. To investigate the involvement of bacterial cysteine proteases single gene mutants of staphopain A and B from the Nebraska transposon mutant library [63] were tested for their cytotoxic potential (Fig 1B). HeLa cells were infected with the wild type strain JE2, the transposon insertion mutants of staphopain A or staphopain B or the non-cytotoxic control strain Cowan I and six hours post-infection LDH release was measured. Whereas the wild type strain showed high levels of cytotoxicity (54.8 ± 8.8 %), infection with the *scp*A transposon mutant led to a significant decrease in cell death (11.9 ± 3.4 %, *p*=0.0011). The plasma membrane damage induced by JE2 was reduced by 78 % by loss of staphopain A function. The *scp*A transposon mutant still exhibited higher cytotoxicity than the control strain Cowan I (−1.3 ± 1.4 %), although cytotoxicity did not differ significantly (*p*=0.1804). By contrast, induction of cell death after infection with the staphopain B transposon mutant (44.2 ± 14.3 %) was slightly, but not significantly decreased in comparison with the wild type (*p*=0.1804) (Fig 1B). The differences in cytotoxicity of the transposon mutants were also observed by morphological analysis of infected HeLa cells thereby corroborating our findings (Fig 1C). Whereas JE2- and JE2 *ssp*B-infected cells showed typical signs of cell death such as cell rounding, retraction of pseudopodia and detachment from the substratum six hours post-infection [64], this phenotype was not observed for cells infected with JE2 *scp*A. There, host cells remained largely adherent, but still had a different appearance when compared to Cowan I infected cells, which looked healthy. On closer examination, most cells infected with the *scp*A transposon mutant were filled with bacteria (Fig 1C). These results suggest that the *S. aureus* cysteine protease staphopain A, but not staphopain B, is involved in cytotoxicity of *S. aureus*. Next, we monitored *S. aureus*-induced cytotoxicity over time. LDH release of HeLa cells infected with wild type bacteria (JE2), *scp*A mutant, *ssp*B mutant or Cowan I was measured over a period of 10 hours (Fig 1D). This assay revealed a delayed onset of cytotoxicity for the *scp*A mutant, which induced significantly less host cell death at 4.5, 6 and 8 hours post-infection when compared to wild type- or JE2 *ssp*B-infected cells. Ten hours post-infection all strains showed the same rate of cytotoxicity. Infection with the staphopain B mutant resulted in slightly, but not significantly reduced intracellular cytotoxicity.

**Fig 1.**
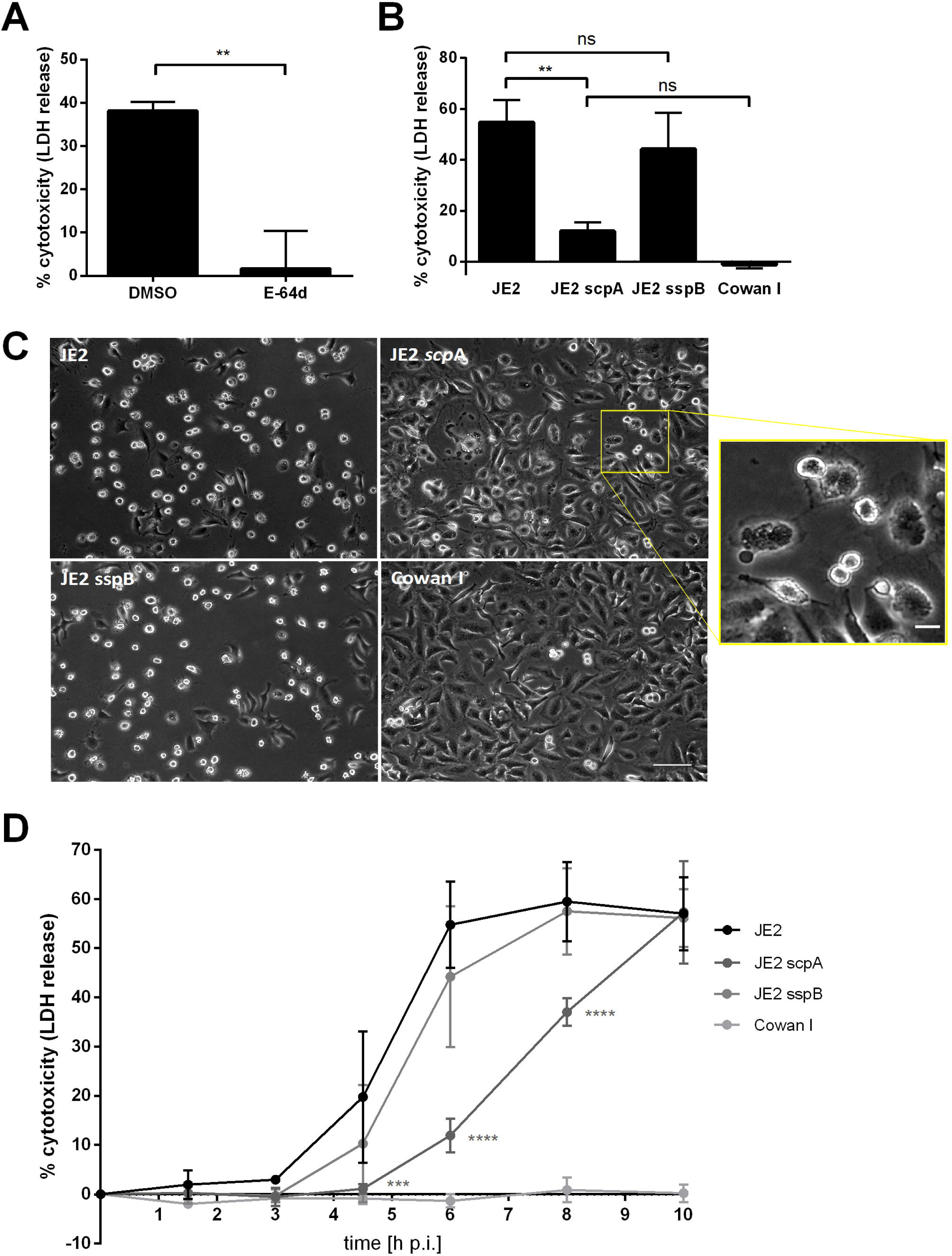
*S. aureus* cysteine protease staphopain A, but not staphopain B, participates in host cell death induced by intracellular bacteria. (A) HeLa cells were treated with 80 µM E-64d or solvent control (DMSO) 1 h prior to infection with *S. aureus* JE2. LDH release of solvent-treated or infected cells was measured 6 h later. (B) HeLa cells were infected with *S. aureus* strains JE2 wild type, staphopain A (JE2 *scp*A) or staphopain B mutant (JE2 *ssp*B) or the non-cytotoxic strain Cowan I and cell death was assessed at 6 h p.i. by LDH release. (C) Phase contrast images of infected HeLa cells at 6 h p.i. revealed different morphologies of cells infected with the *scp*A mutant compared to wild type and *ssp*B mutant (scale bar: 100 µm, scale bar inlet: 20 µm). (D) HeLa cells were infected with wild type strain (JE2), staphopain A mutant (JE2 *scp*A), staphopain B mutant (JE2 *ssp*B) or Cowan I and cell death was assessed at 1.5, 3, 4.5, 6, 8 and 10 h p.i. by LDH assay. Statistical significance was determined by unpaired t test (A), one-way ANOVA (B) or two-way ANOVA (D) (**P<0.01, ****P<0.0001).

To exclude secondary site mutations that interfere with intracellular cytotoxicity of *S. aureus* the transposon inserted in the *scp*A gene was freshly transduced into the JE2 wild type strain and also into the strongly cytotoxic methicillin-sensitive *S. aureus* (MSSA) strain 6850. Infection of HeLa cells with the validated staphopain A mutants exhibited reduced cytotoxicity (JE2 *scp*A: 1.9 ± 1.4 %, 6850 *scp*A: 4.7± 5.5 %) when compared to the wild type six hours post-infection (JE2: 28.8 ± 2.9 %, 6850: 31.6 ± 9.0 %) (Fig 2A, S1A Fig). Loss of staphopain A function led to 93 % reduction of cytotoxicity of *S. aureus* JE2 and 82 % reduction of cytotoxicity of *S. aureus* 6850.

**Fig 2.**
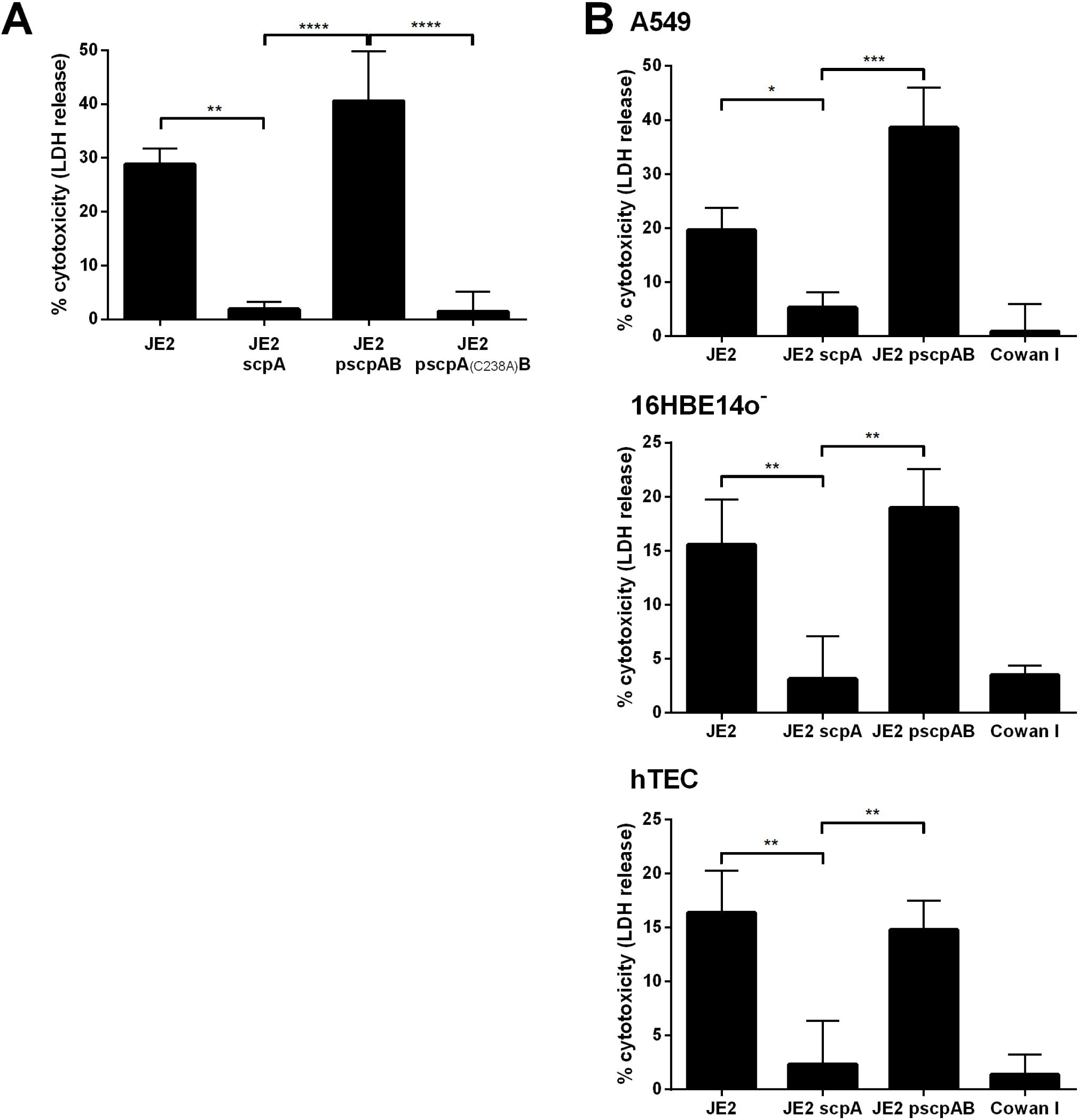
Loss of staphopain A reduces intracellular cytotoxicity of *S. aureus* in different epithelial cell lines and primary cells. (A) HeLa cells were infected with wild type strain (JE2), staphopain A mutant (JE2 *scp*A), complemented mutant (JE2 p*scp*AB) or complemented mutant with active site mutated ScpA (JE2 *scp*A_(C238A)_B) and LDH release was measured 6 h p.i.. (B) Further epithelial cell lines/cells such as A549, 16HBE14o^−^ and human primary tracheal epithelial cells (hTEC) were infected with JE2 wild type, staphopain A mutant (JE2 *scp*A), staphopain B mutant (JE2 *ssp*B) or Cowan I and cell death was assessed at 6 (A549, 16HBE14o^−^) or 8 h p.i. (hTEC) by LDH assay. Statistical significance was determined by one-way ANOVA (*P<0.05, **P<0.01, ***P<0.001, ****P<0.0001).

In order to test whether the less cytotoxic phenotype of the staphopain A mutant can be rescued, the mutant was complemented with p*scp*AB, a plasmid expressing both staphopain A and staphostatin A under the control of their endogenous promotor. As control, the cysteine in the active site of the staphopain A protease domain was substituted by an alanine (C238A) to generate an inactive protease [49]. The *scp*A mutant containing the complementation plasmid induced significantly more LDH release than the uncomplemented mutant in JE2- or 6580-infected HeLa cells, yielding cytotoxicity levels resembling that of wild type *S. aureus* (JE2 p*scp*AB: 40.6 ± 9.2 %, 6850 p*scp*AB: 40.4 ± 5.4 %) (Fig 2A, S1A Fig). Cell death induced by the complemented mutant was even higher compared to the wild type, although not significantly (JE2: *p*=0.0936, 6850: *p*=0.4296), which is concomitant with a higher expression of staphopain A by the high-copy plasmid. By contrast, infection with JE2 p*scp*A_(C238A)_B expressing the *scp*A active site mutant resulted in a low cell death rate (1.4 ± 3.6 %) reminiscent of the *scp*A mutant (Fig 2A). Further, the same kinetics of delayed cytotoxicity were observed for the active site mutant and the staphopain A mutant, while the complemented mutant behaved similarly when compared to the wild type (S1B Fig).

To further investigate the role of staphopain A in *S. aureus*-induced host cell death we performed an apoptosis detection assay by co-staining with annexin V-APC and a cell impermeable, DNA intercalating dye, 7AAD [65]. Wild type, *scp*A mutant or Cowan I-infected HeLa cells were stained with annexin V-APC and 7AAD and subsequent flow cytometrical analysis confirmed that the inactivation of ScpA resulted in reduced host cell death (S1C and D Fig). Infection with the staphopain A mutant led to significantly fewer annexin V**^+^**/7AAD**^+^** cells six hours post-infection as when compared to the wild type, whereas no difference in annexin V**^+^**/7AAD**^+^** cells could be detected between infection with *scp*A mutant and Cowan I. The amount of annexin V**^+^**/7AAD**^−^** and annexin V**^−^**/7AAD**^+^** cells did not reveal any difference between infection with JE2, the *scp*A mutant or Cowan I. This observation indicates a late apoptotic or necrotic behavior of JE2 infected cells at six hours post-infection.

Additionally, we investigated the intracellular effect of staphopain A in other epithelial cell lines (Fig 2B). Adenocarcinomic human alveolar basal cells (A549) and immortalized human bronchial cells (16HBE14o^−^) were infected with *S. aureus* JE2, JE2 *scp*A, JE2 p*scp*AB or Cowan I and six hours post-infection LDH release was quantified. In both cell lines cytotoxicity of the *scp*A mutant was significantly (A549: *p*=0.0371, 16HBE14o^−^: *p*=0.0092) reduced by 73 % or 80 %, respectively, when compared to wild type. Infection with the complemented mutant could restore cytotoxicity in both cell lines. The same results were obtained in primary human tracheal epithelial cells (hTEC) (Fig 2B). Even primary cell infection with JE2 *scp*A led to significantly (*p*=0.0031) less host cell death at early time points of infection, i.e. 8 h p.i.. Mutation of staphopain A reduced cytotoxicity of intracellular *S. aureus* by 86 %, whereas infection with the complemented mutant resulted in cell death rates comparable to the wildtype. Phase contrast and fluorescence microscopy further illustrated these findings (S2 Fig). Epithelial cells infected with the wild type showed cell contraction at six and eight hours post-infection, respectively, whereas infection with JE2 *scp*A did not induce morphological cell alterations, although more intracellular bacteria were observed when compared to Cowan I-infected cells.

### Staphopain A inactivation leads to increased intracellular replication of *S. aureus*

In order to exclude that reduced invasion of the *scp*A mutant accounts for the observed differences in cytotoxicity, we next investigated a role for staphopain A in internalization by the host cell. HeLa cells were infected with wild type bacteria or staphopain A mutant expressing GFP. One hour post-infection, extracellular bacteria were removed by lysostaphin treatment and the percentage of infected cells was determined by flow cytometry (Fig 3A). No significant differences in the amount of host cells infected by either JE2 or JE2 *scp*A were detected (*p*=0.8823). Similarly, CFU counts showed no significant differences in invasion of wild type and *scp*A mutant in HeLa cells (S3A Fig). Therefore, differential internalization does not account for the observed differences in cytotoxicity.

**Fig 3.**
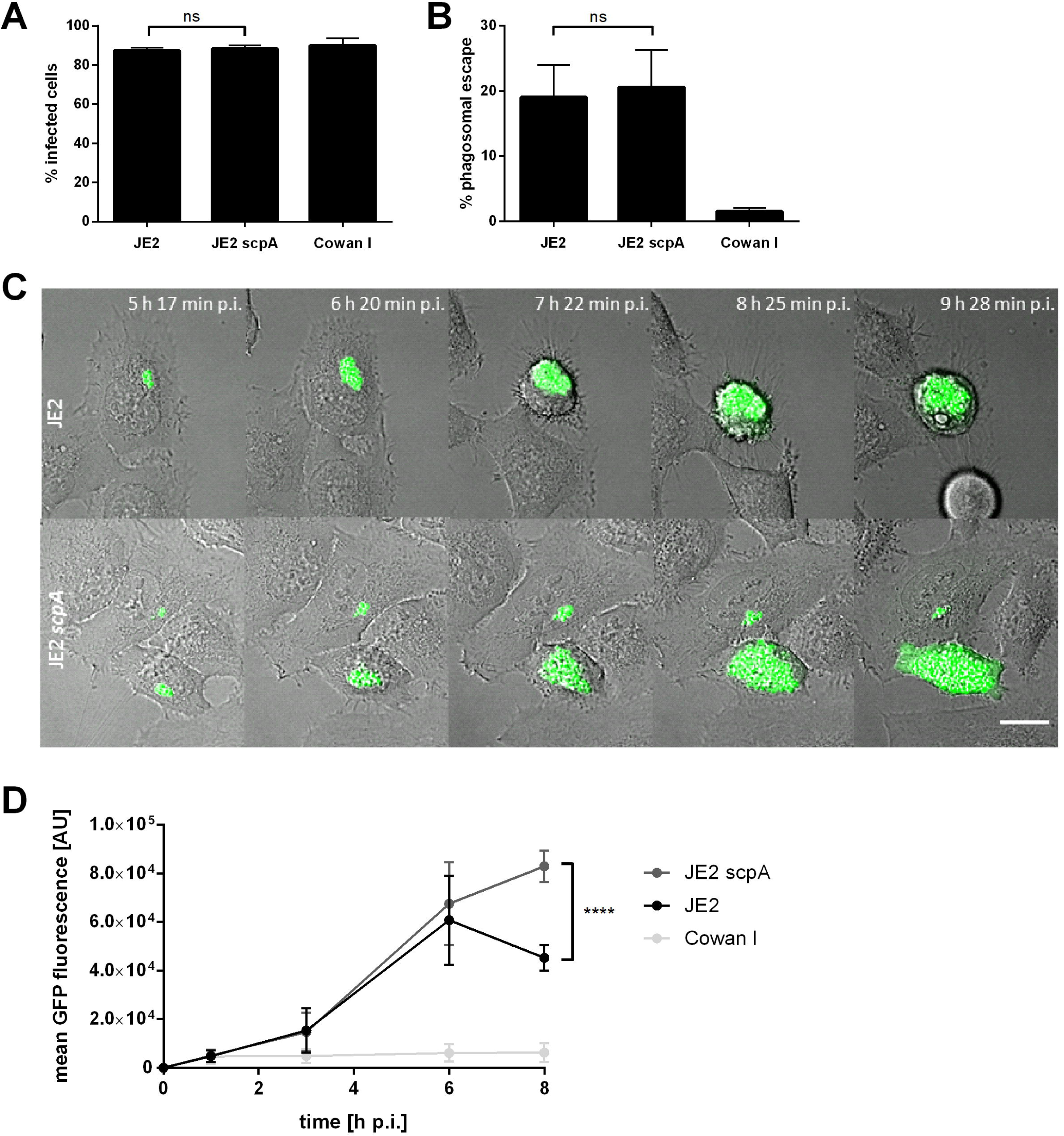
Staphopain A does not interfere with *S. aureus* invasion and phagosomal escape, but prevents excessive intracellular replication in the host cell cytosol. HeLa cells were infected with wild type bacteria (JE2), staphopain A mutant (JE2 *scp*A) or non-cytotoxic Cowan I expressing the fluorescent proteins GFP (A, C, D) or mRFP (B). (A) Invasion into host cells was assessed by measuring the percentage of infected, i.e. GFP-positive, cells at 1 h p.i. by flow cytometry. (B) Phagosomal escape was quantified in the marker cell line HeLa YFP-CWT 3 h p.i. by automated microscopy. (C) Live-cell imaging was performed to study intracellular replication of wild type JE2 (upper panel) and staphopain A mutant (lower panel) by fluorescence microscopy (green: *S. aureus*, gray: BF, scale bar: 20 µm). (D) The mean fluorescence of infected cells was determined at 1, 3, 6 and 8 h p.i. by flow cytometry to analyze intracellular replication. Statistical significance was determined by one-way ANOVA (A, B) or two-way ANOVA (D) (****P<0.0001).

To investigate the role of staphopain A in phagosomal escape, HeLa cells expressing a fluorescent phagosomal escape reporter, YFP-CWT [10, 66], were infected with JE2 wild type and *scp*A mutant and phagosomal escape was recorded by fluorescence microscopy (Fig 3B, S3C Fig, S1 movie). The escape rate of both strains did not show significant differences (*p*=0.8747). Similar results were obtained in 16HBE14o^−^ cells (S3B and D Fig), since both strains escaped from the phagosomes with similar efficieny. Thus, staphopain A is dispensable for *S. aureus* phagosomal escape in epithelial cells.

Escape from the phagosome is followed by intracellular replication of *S. aureus* in the host cell cytoplasm of non-phagocytic cells (S3C and D Fig, S1 movie) [18, 66]. Using live cell imaging we monitored replication of JE2 in HeLa cells, which was accompanied by contraction and rounding of the host cell (Fig 3C, upper panel, S2 movie). The *scp*A mutant displayed substantial intracellular replication, but morphological changes in the infected host cells such as loss of adherence were only observed later during infection (Fig 3C, lower panel, S2 movie). This phenotype of cells infected with the *scp*A mutant was reminiscent of phase contrast images (Fig 1C). Quantification of intracellular replication of *S. aureus* was performed by flow cytometric measurements of bacterial GFP fluorescence and revealed similar rates of JE2 and JE2 *scp*A replication up to six hours post-infection (Fig 3D). Subsequently, the *scp*A mutant continued replicating, whereas the amount of intracellular JE2 dropped as a consequence of their release from lysed cells and killing by extracellular antibiotic. Eight hours post-infection we detected a significant difference in the number of intracellular wild type and mutant bacteria (JE2: 45282 ± 5230 AFU, JE2 *scp*A: 82971 ± 6478 AFU). This correlates with the cytotoxicity of JE2 wild type and staphopain A mutant and suggests that the decrease of intracellular replication of the wild type is caused by a release of the bacteria to the medium, whereas the *scp*A mutant is less cytotoxic and therefore able to replicate longer. CFUs counts of intracellular bacteria further proved these findings (S3E Fig).

### Expression of staphopain A by non-cytotoxic *S. aureus* induces cytotoxicity

We next investigated if expression of staphopain A in an otherwise non-cytotoxic *S. aureus* strain is sufficient to kill the host cell. Non-cytotoxic strains such as Cowan I and the laboratory strain RN4220 are not capable to escape from the phagosome (Fig 3B) [67]. Anhydrous tetracycline (AHT)-inducible expression of δ-toxin (*hld*) has previously been shown to permit phagosomal escape of *S. aureus* RN4220 in human cells, which did not cause a reduction in host cell numbers over a 24-hour period [67]. We therefore generated a transgenic strain of RN4220, which allowed for the AHT-inducible, collinear expression of δ-toxin, staphopain A, staphostatin A and the cyan-fluorescent reporter protein Cerulean. Further, we generated a variant of this plasmid, in which *scp*A was replaced with *scp*A_(C238A)_, harboring a Cys→Ala amino acid substitution rendering the protease inactive [49]. HeLa YFP-CWT cells were infected with recombinant *S. aureus* constructs, either harboring p*hld*-*scp*AB-cerulean or p*hld*-*scp*A_(C238A)_B-cerulean, in the presence of AHT (Fig 4A, S3 movie). Time-lapse microscopy demonstrated phagosomal escape of both strains, which is evidenced by accumulation of the cell-wall binding fluorescence escape reporter around the bacteria (Fig 4A, arrows). HeLa cells infected with the staphopain active site mutant RN4220 p*hld*-*scp*A_(C238A)_B remained adherent and intact, whereas cells infected with RN4220 p*hld*-*scp*AB often were strongly contracted (S4A Fig). Further, only infected host cells, in which *S. aureus* had escaped from the phagosome, showed this contracted phenotype (Fig 4A). Morphological changes induced by cytosolic *S. aureus* expressing functional staphopain A started with the retraction of pseudopodia, followed by strong contraction of the cell and the formation of membrane protrusions (Fig 4B). These morphological changes were similar to the ones observed in HeLa infected with *S. aureus* JE2 (compare Figs 3C and 4B, S1 and S2 movies). However, in HeLa cells infected with the RN4220 p*hld*-*scp*AB the effects were observed within minutes after phagosomal escape, whereas cell shape in JE2-infected HeLa changed only after hours. These differences most likely result from the immediate expression of staphopain A by the transgene, RN4220 p*hld*-*scp*AB. Interestingly, intracellular replication of RN4220 p*hld*-*scp*AB was not needed to induce host cell death (S3 movie). These data suggest that intracellular replication as well as δ-toxin expression is not responsible for the observed morphological host cell changes after escape, but that expression of staphopain A by *S. aureus* in the host cytoplasm leads to cell rounding and subsequent loss of adherence.

**Fig 4.**
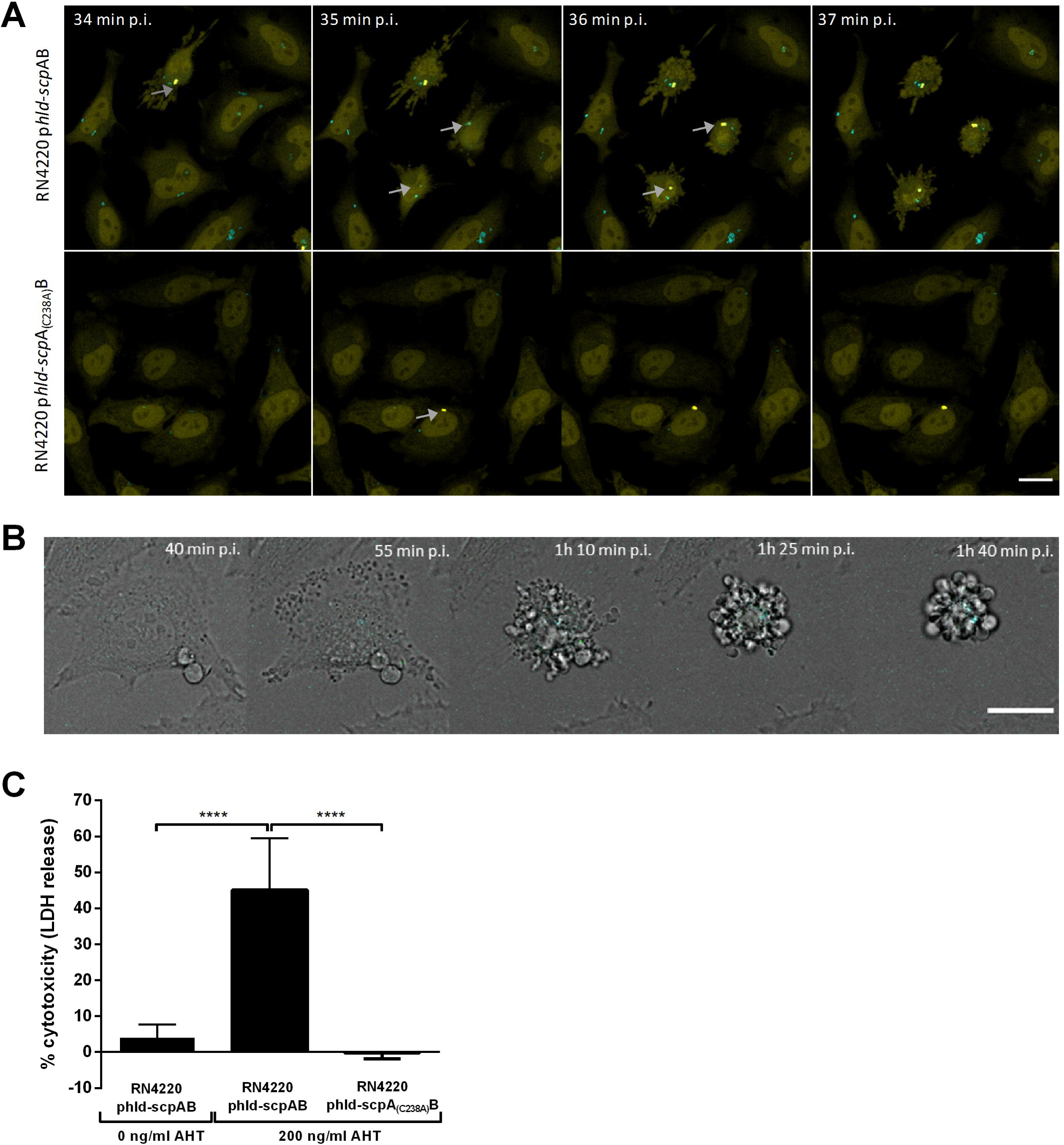
Ectopic expression of staphopain A in a non-cytotoxic *S. aureus* strain induces host cell death after phagosomal escape. (A) HeLa YFP-CWT cells were infected with non-cytotoxic *S. aureus* RN4220 p*hld*-*scp*AB (upper panel) or RN4220 p*hld*-*scp*A_(C238A)_B (lower panel). Live-cell imaging displayed phagosomal escape (see arrows), upon which cell contraction only occurred, when a functional staphopain A was expressed (cyan: *S. aureus*, yellow: YFP-CWT, scale bar: 20 µm). (B) Imaging of HeLa cells infected with *S. aureus* RN4220 p*hld*-*scp*AB visualized cell contraction over time (cyan: *S. aureus*, gray: BF, scale bar: 20 µm). (C) LDH release of HeLa cells was quantified 6 h after infection with *S. aureus* RN4220 p*hld*-*scp*AB or RN4220 p*hld*-*scp*A_(C238A)_B. If indicated, 200 ng/ml AHT was added 1 h prior to infection. Overexpression of functional ScpA resulted in increased plasma membrane damage (n=6). Statistical significance was determined by one-way ANOVA (****P<0.0001).

To elucidate whether the morphological changes induced by escaped RN4220 p*hld*-*scp*AB result in host cell death, an LDH assay was performed six hours post-infection (Fig 4C). Infection of HeLa cells with *S. aureus* RN4220 p*hld*-*scp*AB caused plasma membrane damage when toxin expression was induced by AHT (45.1 ± 14.3 %). By contrast, cytotoxicity of *S. aureus* RN4220 p*hld*-*scp*A_(C238A)_B was absent in the presence of AHT despite induction of δ-toxin (−0.2 ± 1.6 %). In addition, inhibition of protease activity with E-64d in RN4220 p*hld*-*scp*AB infected HeLa cells with AHT treatment significantly reduced LDH release (*p*=0.0010) (S4B Fig). Moreover, RN4220 p*hld*-*scp*AB was significantly less cytotoxic towards HeLa cells when expression was not induced by AHT (4.0 ± 3.6 %) compared to AHT treatment. Cytotoxicity measurements using annexin V- and 7AAD-staining further supported the role of staphopain A in host cell killing (S4C Fig).

In order to exclude a role of extracellular staphopain A in host cell death, *S. aureus* RN4220 p*hld*-*scp*AB and RN4220 p*hld*-*scp*A_(C238A)_B were grown overnight in rich medium supplemented with AHT and supernatant from those cultures was sterile-filtered. The mature and active form of staphopain A was detected in the supernatant of RN4220 p*hld*-*scp*AB, whereas the supernatant of RN4220 p*hld*-*scp*A_(C238A)_B contained only the inactive pro-form but not the mature form of staphopain A (S5A and B Fig). 1, 2 or 5 % of supernatant were added onto fresh HeLa cells and an LDH assay was performed 24 hours after treatment to determine the rate of cell death (Fig 5A). Cytotoxicity of the supernatant was observed in a concentration-dependent manner, but LDH release was comparable in both supernatants, suggesting that virulence factors other than staphopain A were responsible for cell death. In addition, HeLa cells were treated purified staphopain A using various amounts (Fig 5B), but no cytotoxicity was detected. The activity of the purified enzyme was proven (S5C Fig).

**Fig 5.**
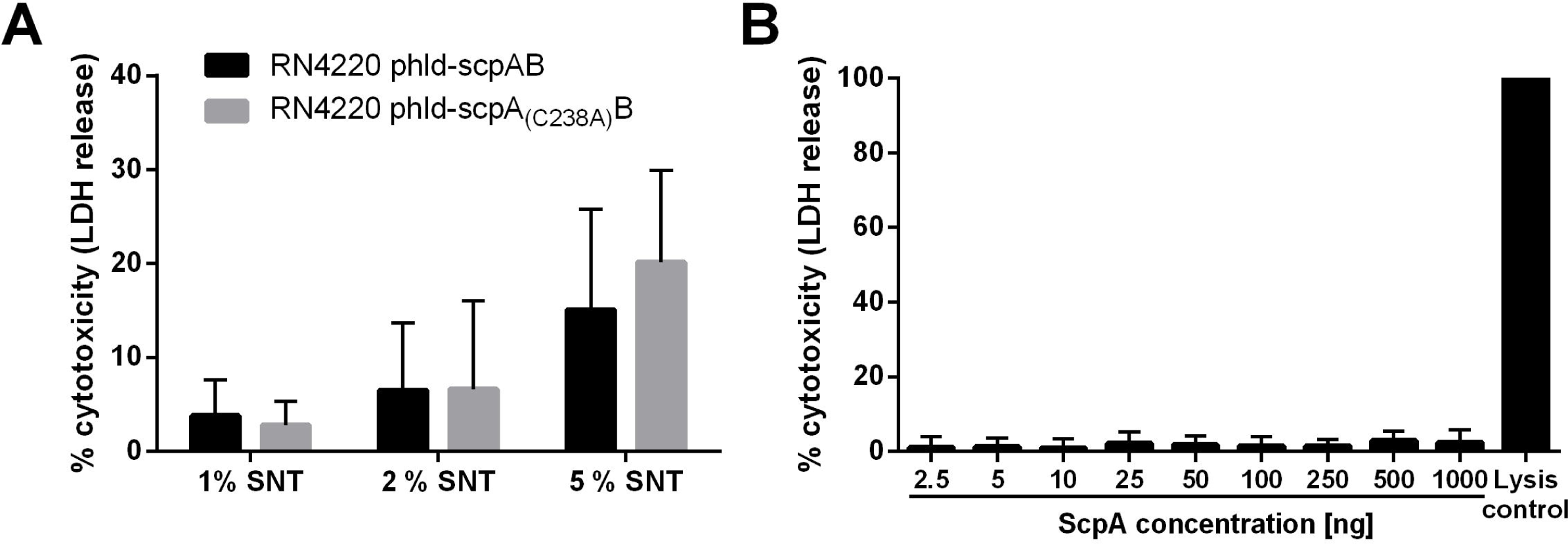
Extracellular staphopain A induces no cytotoxicity. (A) HeLa cells were treated with 1, 2 or 5 % of sterile culture supernatant (SNT) of *S. aureus* RN4220 p*hld*-*scp*AB or RN4220 p*hld*-*scp*A_(C238A)_B or bacterial culture medium as control and 24 h after treatment LDH release was determined. (B) Treatment of HeLa cells with increasing concentrations of purified staphopain A showed no cytotoxicity. Statistical significance was determined by one-way (B) or two-way ANOVA (A).

### Staphopain A-induced cell death displays features of apoptosis

As staphopain A-induced cell death shares similar morphological features with apoptosis like rounding-up of the cell, retraction of pseudopods and plasma membrane blebbing (Fig 4B) [64], we investigated if also molecular characteristics of apoptosis can be found. Annexin V- and 7AAD-staining was performed with RN4220 p*hld*-*scp*AB infected HeLa cells (Fig 6A, S6 Fig). First, infected cells became annexin V-positive followed by additional staining with 7AAD indicating a behavior typical for apoptotic cells. Additionally, apoptosis is usually characterized by the presence of proteolytically active caspases [68]. To investigate the involvement of caspases, HeLa cells were treated with the pan-caspase inhibitor Z-VAD-fmk, infected with *S. aureus* RN4220 p*hld*-*scp*AB and cytotoxicity was determined six hours post-infection by measurement of LDH release (Fig 6B). Inhibition of caspases led to significantly reduced plasma membrane damage induced by infection with RN4220 p*hld*-*scp*AB when compared to untreated cells (*p*=0.0361). However, Z-VAD-fmk was also able to inhibit the activity of purified staphopain A, although not as strongly as the cysteine protease inhibitor E-64 (S6B Fig). Moreover, time-lapse imaging revealed formation of extracellular vesicles at later time points of infection of HeLa cells with RN4220 p*hld*-*scp*AB (Fig 6C, S4 movie). Activity of effector caspases was observed in those vesicles as the fluorescence of a fluorogenic substrate for activated caspase 3/7 increased.

**Fig 6.**
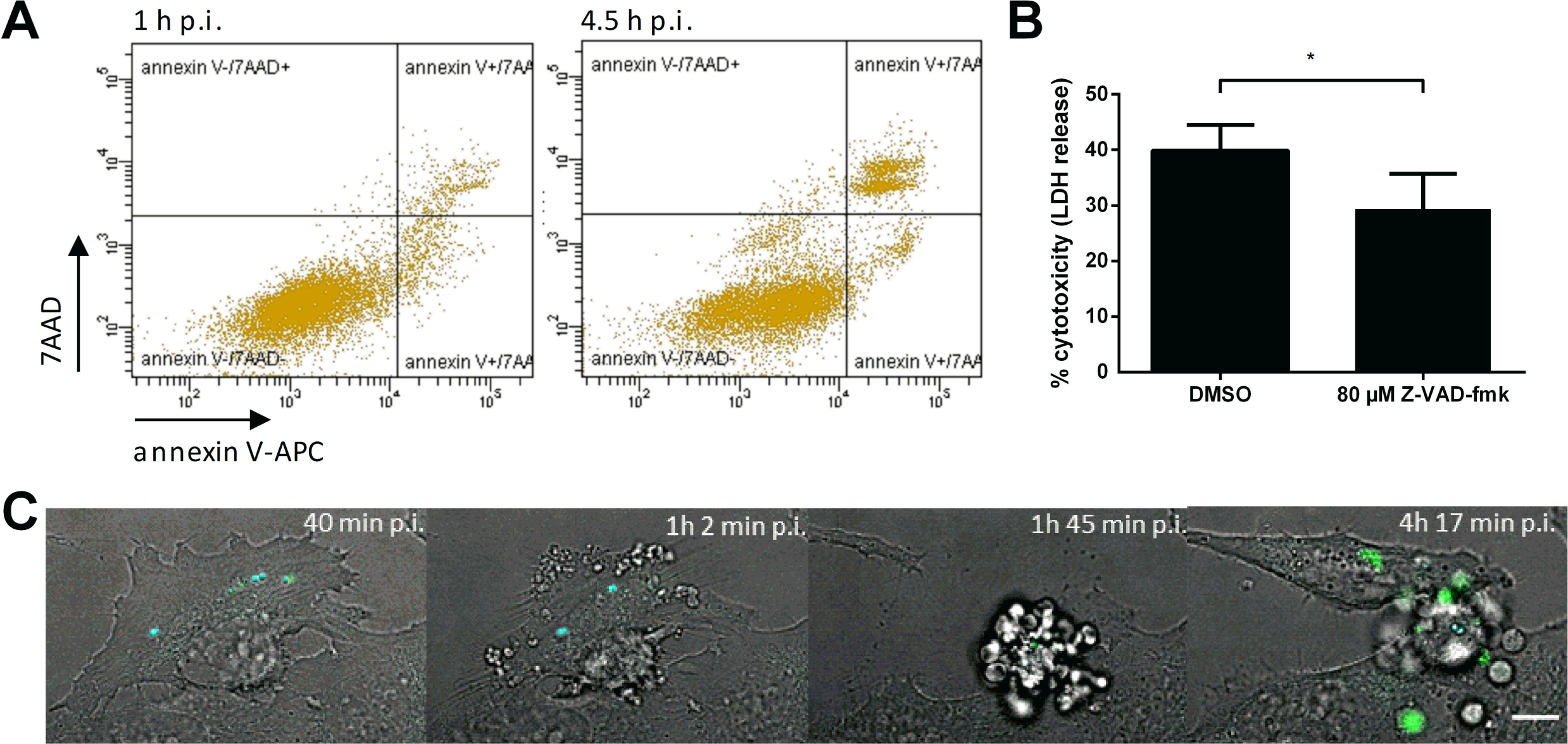
Staphopain A induces apoptotic cell death. HeLa cells were infected with *S. aureus* RN4220 p*hld*-*scp*AB in the presence of 200 ng/ml AHT. (A) Staining with annexin V-APC and 7AAD 1 and 4.5 h p.i. and analysis by flow cytometry to uncover apoptotic cell death of the infected cells. (B) Effect of Z-VAD-fmk (80 µM) on infection-induced release of LDH 6 h p.i. was compared to solvent control (DMSO). (C) Live-cell imaging of infected HeLa cells was performed to monitor the morphology and activation of effector caspases 3/7 (cyan: *S. aureus*, green: CellEvent^TM^ Caspase3/7 Green Detection Reagent, gray: BF, scale bar: 10 µm). Statistical significance was determined by unpaired t test (*P<0.05).

### Inactivation of staphopain A leads to reduced bacterial load in murine lungs

We next tested, if staphopain A was required as a virulence factor in a murine pneumonia infection model. Mice were intranasally infected with equal doses of JE2 wild type, *scp*A mutant or a *scp*A complemented mutant strain. 48 hours after infection mice were sacrificed and bacterial CFUs from the lungs were determined by plating serial dilutions of the tissue lysate (Fig 7). Similar amounts of JE2 and JE2 p*scp*AB were recovered from the lungs, whereas the bacterial load of the *scp*A mutant was significantly reduced (*p*=0.0162). Growth defects of JE2 *scp*A mutant could be excluded (S7A Fig) as well as differential secretion of α-toxin (S7B Fig), which represents the major virulence factor in *S. aureus* induced pneumonia [69]. Thus, insertional inactivation of staphopain A resulted in attenuated virulence of *S. aureus* JE2 in murine lungs.

**Fig 7.**
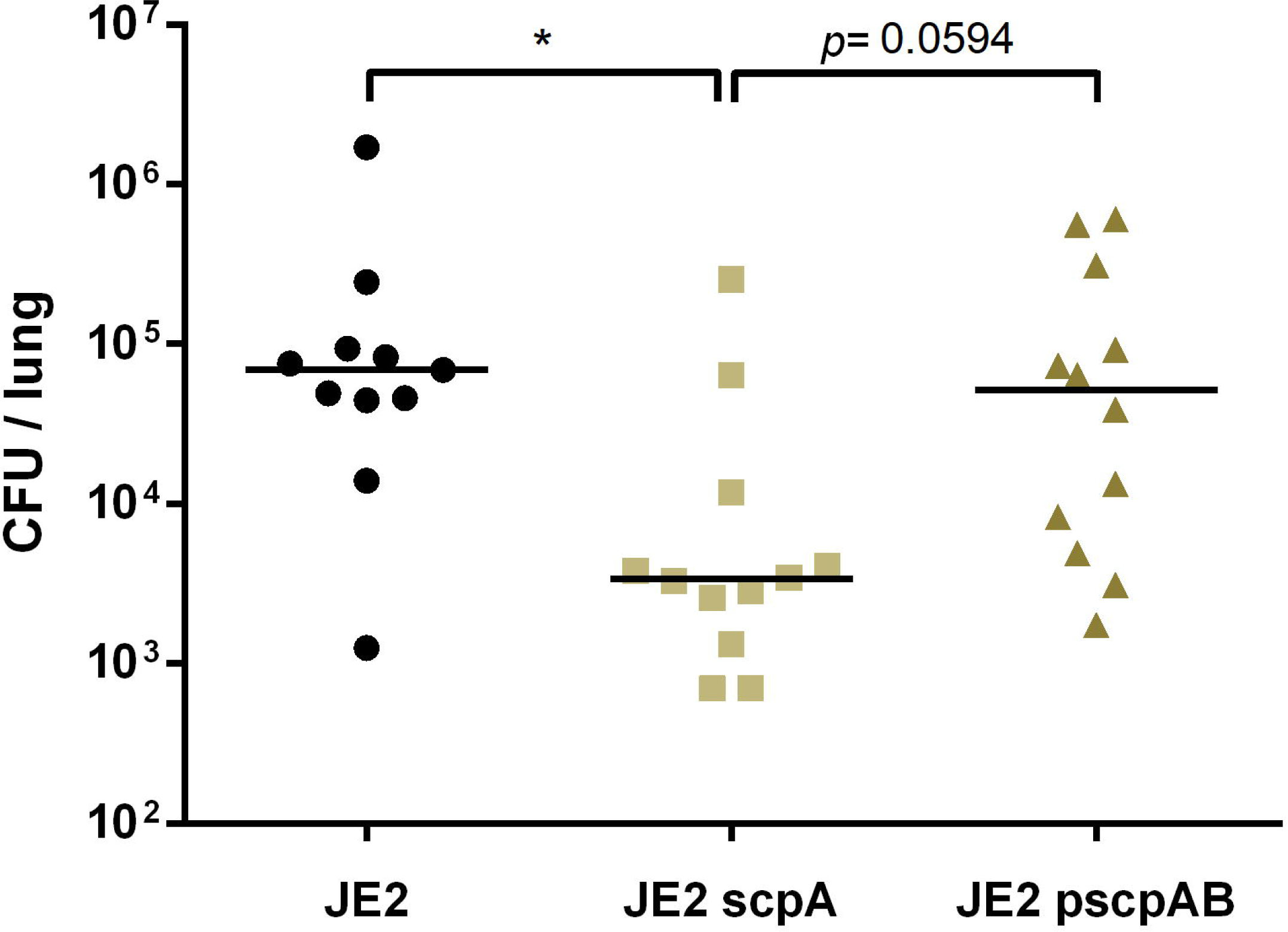
*In vivo* effects of staphopain A loss of function. Balb/c mice were intranasally administered with wild type bacteria (JE2), *scp*A mutant (JE2 *scp*A) or complemented mutant (JE2 p*scp*AB) and bacterial CFUs were recovered from lung tissue 48 h p.i. by plating serial dilutions of the lysed tissue. The horizontal line represents the median of recovered CFUs from total lungs of mice, individual points illustrate the CFUs recovered from the lungs of one mouse. Statistical significance was determined by Kruskal-Wallis test (*P<0.05).

## Discussion

*S. aureus* is able to invade epithelial and endothelial cells and causes cell death after phagosomal escape. Whereas phenol soluble modulins are important in translocation of endocytosed bacteria to the host cytosol [66], not much is known on the downstream processes and the respective virulence factors involved. Aside from pore-forming toxins *S. aureus* secretes a multitude of virulence factors such as lipases and proteases. It has been reported that cysteine proteases facilitate escape of intracellular pathogens from the host cell. For instance, a papain-like cysteine protease of the malarial parasite *Plasmodium falciparum* has been shown to be essential for rupture of the host cell membrane [70]. Furthermore, cysteine proteases of mammalian cells like caspases, calpains and cathepsins are associated with cell death [71]. Of the two cysteine proteases encoded by *S. aureus*, only staphopain B has been connected to *S. aureus* pathogenicity, whereas the virulence potential of staphopain A is disputed, since the mutant showed no effect in animal studies [72, 73]. However, it was shown that staphopain A, for instance, cleaves the chemokine receptor CXCR2 *in vitro* thereby blocking neutrophil activation and chemotaxis [53]. Here, we investigated the contribution of staphopain A and staphopain B to *S. aureus*-induced intracellular cytotoxicity. Treatment with the cysteine protease inhibitor E-64d prior to infection led to a significant reduction in death of host cells infected with *S. aureus* (Fig 1A) thereby suggesting an involvement of cysteine proteases of either host or pathogen origin in the pathogen-induced host cell death. Inhibition of cysteine proteases by E-64 in a murine ocular infection model also resulted in reduced *S. aureus* virulence, as shown by reduced ocular injury and enhanced bacterial clearance [74]. We therefore tested insertional *S. aureus* mutants within either of the structural genes, *scp*A and *ssp*B, for cell death phenotypes. Interestingly, only the loss of staphopain A led to drastic reduction of cytotoxicity of both tested *S. aureus* strains, JE2 and 6850 (Fig 1B and D, S1A Fig). The loss of cytotoxicity in the *scp*A mutant was readily complemented in trans by reintroduction of a functional *scp*A ORF under control of its native promoter, but was absent from an active site mutant in *scp*A (C238A) [49] (Fig 2A, S1B Fig). Involvement of staphopain A in intracellular cytotoxicity of *S. aureus* was not only detected in cancer cell lines, but also in immortalized and even primary epithelial cells (Fig 2B). Our data thus demonstrate a novel role of ScpA during intracellular *S. aureus* pathogenesis, which is dependent on the proteolytic activity of staphopain A.

Bacterial extracellular proteases, such as staphopain A, process bacterial cell surface proteins, host cell receptors or extracellular matrix proteins, which are crucial for bacterial invasion [75]. Also, enhanced *S. aureus* cytotoxicity is associated with increased host cell invasion [8]. However, staphopain A neither was involved in host cell invasion nor in phagosomal escape (Fig 3A and B, S3A-D Fig, S1 movie), another process that is implicated in intracellular *S. aureus* cytotoxicity in non-professional phagocytes [10, 66, 76].

Loss of staphopain A function led to a delayed onset of host cell death induced by intracellular *S. aureus* and permitted a prolonged intracellular residence and replication of the pathogen (Fig 1D, 3C and D). Our group previously identified a loss-of-function mutant of the pleiotropic transcriptional regulator Rsp [77]. Mutants in *rsp* demonstrate a delayed host cell death, which is reminiscent of the data obtained in the present study. Since staphopain A expression was shown to be dependent on Rsp [77], we suggest that the attenuated-cytotoxicity phenotype and the prolonged intracellular residence of the *rsp* mutant is at least partially due to diminished expression of staphopain A. Transcription of *scp*A was reported to be increased in *S. aureus* residing in THP1 macrophages [78].

We next used the laboratory cloning strain *S. aureus* RN4220 and engineered it to allow controlled release of staphopain A into the host cytosol by inducible co-expression of the PSM δ-toxin, as well as *scp*AB and the fluorescent marker Cerulean. Expression of δ-toxin led to translocation of the transgenic bacteria into the host cytosol [67]. Subsequently, the strain expressing wild type *scp*A caused host cell rounding and detachment from the substratum, whereas expression of the active site mutant *scp*A (*scp*A_(C238A)_) did not (Fig 4A and B, S4A Fig). Cell death caused by RN4220 p*hld*-*scp*AB was only detectable, when *S. aureus* escaped into the cytoplasm (Fig 4A and C), highlighting the dependency of *S. aureus* cytotoxicity on phagosomal escape and pointing to an intracellular target substrate of staphopain A in epithelial cells. Whether staphopain A acts alone or in concert with other bacterial factors, cannot be completely answered. Although transgenic RN4220 escaped to the cytoplasm of infected HeLa cells, it did not grow in this environment irrespective of the presence of a functional ScpA protease (S4 movie), illustrating that ScpA is not responsible for intracytoplasmic growth of *S. aureus*. RN4220 exhibits altered expression of virulence factors, for instance this strain shows delayed expression of the major virulence regulator *agr* with small amounts of RNAIII and failure to translate α- and δ-toxin [79], which is based on a mutation in *agr*A [80]. Hence, the *agr* deficiency or other mutations in this cloning strain may cause the inability to replicate in the host cell cytoplasm. Besides, ectopic expression of staphopain A induces cell death rather quickly, which in turn may not allow bacterial replication (Fig 4A).

We excluded extracellular staphopain A to account for cytotoxicity to HeLa cells (Fig 5). Although we observed concentration-dependent cell lysis of sterile culture supernatant from *S. aureus* RN4220 expressing δ-toxin and staphopain A, cell death rates of ScpA and ScpA_(C238A)_ were comparable suggesting that other factors than ScpA such as δ-toxin induce in cytolysis under these conditions [81]. Interestingly, purified staphopain B induced cell death in human neutrophils and monocytes, but not in other cell types, such as macrophages and epithelial and endothelial cells [82]. We observed no cytotoxic effect of purified staphopain A on epithelial cells (Fig 5B).

Inducible expression of *scp*A by *S. aureus* in HeLa cytosol indicated an apoptotic mode of cell death since the host cells first became annexin V positive and only later were stained by 7AAD (Fig 6A). This is a typical representation of apoptosis, where lysis of the plasma membrane is preceded by presentation of phosphatidylserines (PS) on the outer leaflet of the cell membrane [83]. Moreover, cytotoxicity of *scp*A-expressing *S. aureus* was attenuated after treatment with the pan-caspase inhibitor, Z-VAD-fmk (Fig 6B). Further, we observed activity of the effector caspases 3/7 during staphopain A-induced cell death (Fig 6C, S4 movie). Together with morphological characteristics of apoptotic cells, such as cell rounding, retraction of pseudopods, plasma membrane blebbing and formation of extracellular vesicles reminiscent of apoptotic bodies [64, 84], our results suggest that ScpA expressed by *S. aureus* in the cytoplasm of host cells leads to an apoptosis-like cell death. This process may be followed by secondary necrosis [85], as we observed LDH release and 7AAD staining. Interestingly, cathepsin B was identified as the closest structural neighbor of staphopains among the eukaryotic enzymes [48] and the overall substrate preferences of ScpA and SspB most closely resembled that of human cathepsin H [86]. Cathepsins are eukaryotic proteases, which are associated with cellular protein turnover, but also induce apoptosis by cleavage of Bid and degradation of the anti-apoptotic Bcl-2 homologues [87–90]. In addition, the trypanosomal cysteine cathepsin cruzipain directly activates caspases 3/7 [90]. Therefore, it is conceivable that staphopain A induces host cell apoptosis in a similar manner. Staphopain A has also been reported to liberate the activity of cathepsin B *in vitro* by proteolytic degradation of cystatin C, a natural inhibitor of cysteine proteases [91].

However, treatment with Z-VAD-fmk could not fully abolish cytotoxicity of *S. aureus* RN4220 p*hld*-*scp*AB (Fig 6B) and ScpA activity was decreased by pan-caspase inhibitor treatment *in vitro* (Fig S6). Therefore, ScpA may also induce other cell death pathways, besides apoptosis. Further, cytotoxicity experiments using Z-VAD-fmk treating *S. aureus* infection should be carefully evaluated. This finding may also explain the controversies on apoptosis induced by *S. aureus* infection [18].

Host cell death mediated by intracellular *S. aureus* independently of staphopain A may induce mechanical stress to the host cell due to a high number of intracellular bacteria occupying the cytoplasm of the host cell. Host cells infected with JE2 *scp*A appear to be fully packed with bacteria (Fig. 1C and S2, Movie S2). This likely results in a necrotic type of cell death. On the other hand, another *S. aureus* virulence factor may be able to trigger late host cell death in addition to and independently of staphopain A. This factor may require increased bacterial density or a certain critical concentration. Metabolite shortage induced by replicating bacteria in the host cell is yet another possibility of how infection stress could impact cell viability independent of staphopain A.

A *S. aureus* mutant lacking ten of the major extracellular proteases, including staphopain A, showed higher mortality and conversely less bacterial burden in the lungs, liver, heart and spleen, but not in brain and kidneys in a murine sepsis model and exhibited reduced bacterial loads per abscess in a murine model of skin abscess [92]. Interestingly, numbers of intracellular bacteria in professional phagocytes of whole human blood were increased for the protease-null mutant of *S. aureus* compared to the wild type [92], supporting our finding in HeLa cells where loss of staphopain A delays bacterial induced cell death and thus enables prolonged intracellular replication. However, the observed phenotype in the aforementioned study cannot be clearly assigned to staphopain A, since additional proteases were inactivated in the strain used. Additionally, expression of staphopain A was enhanced *in vivo* during infection in a murine osteomyelitis model compared to *in vitro* exponential growth [93]. Here, we employed a murine lung infection model to demonstrate the *in vivo* relevance of *S. aureus* staphopain A and found less bacterial burden in the lung tissue of infected mice, when staphopain A was mutated in comparison to the wild type (Fig 7). We used the highly cytotoxic *S. aureus* USA300 derivative JE2, similarly to Kolar et al. [92], whereas another study, which detected no effect of ScpA loss of function on virulence in a mouse abscess model [72], infected with low cytotoxic *S. aureus* 8325-4 [40]. Hence, the infection models of the latter study may not be suitable to reveal the contribution of staphopain A on cytotoxicity of *S. aureus in vivo*.

Quantification of intracellular *S. aureus* in HeLa cells revealed that loss of staphopain A function led to increased bacterial growth due to prolonged intracellular residence (Fig 3C and 3D, S2 movie). It is conceivable that this longer intracellular residence time is the reason for the poorer colonization efficiency of the staphopain A mutant in the murine lung, as the wild type may kill the host cell more rapidly and thus lead to more extensive tissue destruction *in vivo*. This may allow *S. aureus* to establish infection of the organ more efficiently when compared to the mutant. However, these findings cannot be directly compared, since strictly intracellular bacteria were quantified in the *in vitro* experiment with HeLa cells, whereas the bacterial location was not determined for the *in vivo* experiment. The *in vivo* role of intracellular *S. aureus* therefore requires more extensive investigation. It was shown that *S. aureus* can survive within alveolar macrophages and neutrophils, which are the first line of defense in the lung [20]. Nevertheless, we currently cannot exclude extracellular effects of staphopain A in the murine lung. For instance, a study showed that ScpA is an important virulence factor that can impair innate immunity of the lung through degradation of lung surfactant protein A (SP-A) [94]. Cleavage of the chemokine receptor CXCR2 by staphopain A and thereby blocking of neutrophil recruitment may also play a role in *S. aureus* pathogenicity *in vivo* [53].

In summary, we demonstrate in the present study that *S. aureus* cysteine protease staphopain A induces cell death in epithelial cells after translocation to the host cell cytoplasm. Whereas ScpA is not involved in cell invasion and phagosomal escape, it activates a mode of cell death with hallmarks of apoptosis. Cell death induced by *scp*A mutants is delayed by multiple hours, during which bacteria can replicate in the host cytosol. We hypothesize that intracellular *S. aureus* utilizes staphopain A to exit the host cell thereby leading to tissue destruction and potentially dissemination of infection. In addition, staphopain A is required for efficient colonization of the mouse lung by cytotoxic *S. aureus* suggesting that this protease and its cytotoxic activity may be crucial for the virulence of the pathogen *in vivo*.

## Methods

### Bacterial culture conditions

*Escherichia coli* strains were grown in Luria-Bertani broth (LB) and *Staphylococcus aureus* strains were grown in Tryptic soy broth (TSB, Sigma), if not stated otherwise. Media were supplemented with appropriate antibiotics, when necessary, and broth cultures were grown aerobically at 37°C overnight at 180 rpm. *E. coli* was selected on LB plates containing 100 µg/ml ampicillin and selective TSB plates for *S. aureus* were prepared using 10 µg/ml chloramphenicol and/or 5 µg/ml erythromycin (for chromosomally encoded antibiotic resistance).

### Bacterial growth curves

Bacterial growth curves were measured with a TECAN InfiniteM Plex plate reader. Bacterial cultures were inoculated in triplicates to an OD_600nm_ of 0.1 in 400 µl TSB and grown for 18 h at 37 °C in a 48 well microtiter plate. Absorbance was recorded every 10 minutes at 600 nm.

### Construction of bacterial strains and plasmids

All used strains, plasmids and oligonucleotides can be found in S1-S3 tables in the supplemental material. The *S. aureus* insertional transposon mutant of staphopain A (NE1278) from the Nebraska Transposon mutant library [63] was transduced via phage φ11 into the genetic background of wild-type *S. aureus* JE2 and 6850 in order to avoid secondary site mutations.

For complementation of staphopain A the *scp*AB operon including the native promotor region, 361 bp upstream of start codon, and the transcription termination signal, 253 bp downstream of the stop codon, was amplified by PCR (for primer see S3 table) from genomic DNA of *S. aureus* JE2 and gfp_uvr_ in p2085 was replaced by the generated insert using cloning sites PstI and EcoRI.

Site directed mutagenesis for Cys_238_>Ala active site substitution in *scp*A was performed using the QuikChange II Site-Directed Mutagenesis Kit (Agilent Technologies) using oligonucleotides MP_*scp*A1 and MP_*scp*A2 [49].

Plasmids pmRFPmars and pGFPsf were transduced into the respective strains (see S1 table) using phage φ11. pGFPsf was constructed by cloning sarAP1-mRFPmars from pmRFPmars into the pSK5632 backbone using cloning sites PscI and KasI and replacing mRFPmars with synthetic, codon-adapted Superfolder GFP (GFPsf, GeneArt, ThermoFisher) using cloning sites AvrII and BamHI.

For construction of p*hld*-*scp*AB-cerulean and p*hld*-*scp*A_(C238A)_B-cerulean the *scp*AB operon was amplified from p*scp*AB or p*scp*A_(C238A)_B, respectively, using primers *scp*AB_AvrII_fwd and *scp*AB_AvrII_rev and cloned into p*hld*-*hlb*-cerulean by replacing *hlb* using AvrII restriction sites.

All assembled vectors were transformed into chemically competent *E. coli* DH5α and confirmed via PCR and Sanger sequencing (SeqLab, Göttingen). Subsequently, vectors were electroporated into *S. aureus* RN4220 and, if required, further transduced into *S. aureus* JE2 or 6580 using phage φ11.

### Preparation of sterile supernatant

Bacteria were grown overnight at 180 rpm in BHI medium (Sigma Aldrich) supplemented with antibiotics and 200 ng/µl AHT, if indicated. Cultures were adjusted to an OD_600nm_ of 10. Subsequently, bacterial cultures were centrifuged and the supernatant was sterile filtered (0.22 µm pore size).

### Purification of Staphopain A

*S. aureus* strain V8 (BC10 variant) was grown overnight in TSB (4 l) supplemented β-glycerophosphate (5 mg/mL), bacterial cells were removed by centrifugation (5,000 × g, 4 °C, 20 min) and proteins in supernatant were precipitated at 4°C by slow addition of AmSO_4_ to 80% concentration (561 g/1 l) with continuous steering. After the last portion of AmSO_4_, the sample was stirred for 60 min and then precipitated proteins were pelleted by centrifugation (10,000 × g, 4 °C, 30 min) and re-suspended in ice cold 50 mM sodium acetate, pH 5.5 (NaAc5.5). After extensive dialysis (24 h, against 2 l NaAc5.5 with 3 changes, 4°C) precipitate was removed by centrifugation (20,000 × g, 4 °C, 20 min) and the supernatant passed through 0.45 µm filter. The sample was loaded on a HiPrep Q Fast Flow 16/10 (#28-9365-43, GE Healthcare) column equilibrated with NaAc5.5. The column was washed with NaAc5.5 at a flow rate of 1 ml/min and 8-10 ml fractions were collected. Staphopain A (ScpA) proteolytic activity was determined using azocasein and active fractions were pooled. Of note, staphopain B (SspB), V8 protease and aureolysin bind HiPrep Q FF and could be eluted with NaCl gradient. Next, a pool containing Staphopain A was directly loaded on a HiPrep CM Fast Flow 16/10 (#28-9365-42, GE Healthcare) column equilibrated with NaAc5.5 and the column was washed until OD_280nm_ reached a baseline. Staphopain A was eluted with NaCl gradient from 0 to 0.2 M in the total volume of 500 ml developed at the flow rate of 1 ml/min. Fractions containing proteolytic activity on azocasein in the presence of 2 mM dithiothreitol (DTT) and 5 mM EDTA were pooled and concentrated by ultrafiltration on Amicon 10 kDa cut-of membrane and loaded on HiLoad 16/60 Superdex 75 pg (#28-9893-33, GE Healthcare) equilibrated with NaAc5.5 and fractions containing the proteolytic activity on azocasein were pooled and concentrated as described above. The procedure yielded up to 10 mg of >95% pure staphopain A (ScpA) (by SDS-PAGE) free of other staphylococcal proteases.

### Staphopain A activity assay

The fluorogenic substrate Z-Leu-Leu-Glu-AMC (#S-230-05M, R&D) was used to monitor staphopain A activity *in vitro*. 200 µM substrate was mixed with 50 µl 4x buffer (0.4 M sodium phosphate pH 7.4, 5 mM EDTA, freshly supplemented with 8 mM DTT) and 50 µl sterile bacterial culture supernatant in a total volume of 200 µl and transferred into 96 well plate. The plate was incubated for 3 h at 37 °C and mean fluorescence intensity with emission at 445 nm with excitation at 345 nm was measured with a fluorescence plate reader. For each sample values were subtracted from a blank sample (without enzyme). Activity of purified staphopain A was determined with the same protocol, except plates were incubated for 22 h until fluorescence measurement. To investigate the inhibitory activity of E-64 (Merck) and Z-VAD-fmk (Invivogen) various concentrations of the substances were added to the substrate-buffer mix with 200 ng purified enzyme.

Alternatively, staphopain A activity was determined with azocasein (1% final concentration) as the substrate using 0.1 M Tris, 5 mM EDTA, pH 7.6 freshly supplemented with 2 mM DTT. Briefly, enzyme sample was mixed with assay buffer in the total volume of 200 µl. After temperature was adjusted to 37 °C, 100 µl of 3% azocasein in the assay buffer was added. After 1 h the reaction was stopped by addition of 200 µl of 10 % ice cold trichloroacetic acid (TCA). After 10 min on ice precipitate was removed by centrifugation (10,000-12,000 × g, 5 min, 4 °C), 200 µl were transferred to a microplate and OD_360nm_ was measured against a blank sample containing all regents but the enzyme samples.

### Infection of epithelial cells

HeLa cells (HeLa 229, ATCC CCL-2.1) were grown in RPMI1640 medium (#72400021, ThermoFisher Scientific) and 16HBE14o^−^ [95] and A549 cells were grown in DMEM (D6429, Sigma Aldrich) supplemented with 10 % FBS (Sigma Aldrich) and 1 mM sodium pyruvate (ThermoFisher Scientific) at 37 °C and 5 % CO_2_. For infection 0.8 to 1 × 10^5^ cells were seeded into 12 well microtiter plates 24 hours prior to infection. 1 hour prior to infection medium was renewed and, if required, treatment with 80 µM E-64d (Merck), 80 µM Z-VAD-fmk (Invivogen) or 200 ng/ml anhydrous tetracycline (AHT) was applied.

Bacterial overnight cultures were diluted to an OD_600nm_ of 0.4 and incubated for 1 hour at 37 °C and 180 rpm to reach exponential growth phase. Then, bacteria were washed twice by centrifugation and used to infect HeLa cells at a multiplicity of infection (MOI) of 50, if not stated otherwise. After one hour co-cultivation extracellular bacteria were removed by 30 minutes treatment with 20 µg/ml lysostaphin (AMBI) followed by washing and further incubation in medium containing 2 µg/ml lysostaphin and, if required, 80 µM E-64d, 80 µM Z-VAD-fmk or AHT until the end of experiment.

### Isolation and infection of primary cells

Isolation of primary human tracheal epithelial cells (hTEC) was performed as described previously [96]. Briefly, the airway mucosa was removed mechanically from human tracheobronchial biopsies, which were subsequently placed into plastic cell culture dishes and covered with Airway Epithelial Cell Growth Medium (AECG, # PB-C-MH-350-0099, PeloBiotech). After 8-12 days hTEC grown out of the tissue pieces were collected. Infection of primary cells was performed as described above for epithelial cell lines. Except, 1.75 × 10^5^ cells were seeded into 24 well microtiter plates 24 hours prior to infection.

### LDH assay

HeLa cells were infected as described above and 1.5 hours after infection medium was replaced by RPMI1640 without phenol red containing 1 % FBS and 2 µg/ml lysostaphin. 6 hours after infection medium was removed from the wells, shortly centrifuged and 100 µl of supernatant of each sample were transferred into the well of a 96 well microtiter plate in triplicates. LDH release was measured using the Cytotoxicity Detection Kit Plus (Roche) according to manufacturer’s instruction.

For LDH assay with sterile supernatant, 1, 2 or 5 % dilutions of sterile bacterial supernatant were prepared in RPMI1640 medium containing 1 % FBS and no phenol red and added to fresh HeLa cells. 24 hours after treatment LDH release was measured as described above. Uninfected cells served as negative control and lysed cells served as positive control.

### Annexin V and 7AAD staining

HeLa cells were infected as described above. At the desired time point after infection medium, which possibly contained detached dead cells, was collected from the wells and adherent cells were detached using TrypLE (ThermoFisher Scientific). Adherent and suspension cells of each sample were pooled and after centrifugation for 5min at 800 × g cells were carefully resuspended in annexin V staining buffer (RPMI1640 without phenol red containing 1 % FBS, 2 µg/ml lysostaphin, 2 mM CaCl_2_, 10 µl/ml annexin V-APC [BD Biosciences] and 10 µl/ml 7AAD [BD Biosciences]). After 10 minutes incubation in the dark cells were immediately analyzed by flow cytometry using a FACS Aria III (BD Biosciences) and BD FACSDiva Software (BD Biosciences). Forward and sideward scatter (FSC-A and SSC-A) were used to identify the cell population and doublet discrimination was performed via FSC-H vs. FSC-W and SSC-H vs. SSC-W gating strategy. APC or 7AAD fluorescence was measured using a 633 nm or 561nm laser for excitation and a 660/20 nm or 610/20 nm band pass filter for detection, respectively. 10,000 events were recorded for each sample. Uninfected cells served as negative control.

### Flow cytometry-based invasion assay

HeLa cells were infected with GFP expressing strains and prepared for flow cytometry as described above. Detached cells were resuspended in fresh medium without phenol red containing 1 % FBS and 2 µg/ml lysostaphin one hour post infection, after 10 minutes treatment with 20 µg/ml lysostaphin to remove extracellular bacteria. For determining invasion, the percentage of GFP-positive cells representing the infected cells was measured by flow cytometry using a FACS Aria III (BD Biosciences). Gating and analysis were performed as described above. GFP fluorescence was measured using a 488nm laser and a 530/30 nm band pass filter for detection. Uninfected cells were used to determine autofluorescence of the cells and signals above this value were defined as infected.

### Flow cytometry-based intracellular replication assay

Cells were infected with GFP expressing strains and prepared for flow cytometry as described above. Intracellular replication was determined by measurement of GFP fluorescence (arbitrary units, AU) of the infected cells 1, 3, 6 and 8 hours after infection by flow cytometry using a FACS Aria III (BD Biosciences). The intensity of GFP fluorescence corresponds to the amount of intracellular bacteria. Gating and analysis were performed as described above. GFP fluorescence was measured using a 488nm laser and a 530/30 nm band pass filter for detection. Uninfected cells served as negative control.

Bacterial invasion into the host cells as well as intracellular replication were additionally analyzed by counting bacterial colony forming units (CFU) recovered from infected cells.

### Phagosomal escape assay

Phagosomal escape was determined as described previously with minor modifications [10]. Briefly, HeLa YFP-CWT cells were infected with mRFP-expressing bacterial strains at a MOI of 10 in a 24 well µ-plate (ibidi). After synchronization of infection by centrifugation, infected cells were incubated for 1 hour to allow bacterial invasion. Subsequently, a 30 minute-treatment with 20 µg/ml lysostaphin removed extracellular bacteria, after which the cells were washed and medium with 2 µg/ml lysostaphin was added. Three hours after infection, cells were washed, fixed with 4 % paraformaldehyde overnight at 4 °C, permeabilized with 0.1% Triton X-100 and nuclei were stained with Hoechst 34580. Images were acquired with an Operetta automated microscopy system (Perkin-Elmer) and analyzed with the included Harmony Software. Co-localization of YFP-CWT and mRFP signals indicated phagosomal escape.

### CFU assay

Infection was performed as described above. After one hour bacteria-host cell co-cultivation, extracellular bacteria were removed by treatment with 20 µg/ml lysostaphin.

Bacterial invasion was assessed by employing a 10-minute lysostaphin treatment, followed by washing with sterile PBS and subsequent osmotic shock-mediated lysis with alkaline water (pH 11) for 5 minutes at room temperature. Dilution series of the lysate were plated on tryptic soy agar (TSA) and incubated overnight at 37 °C to enumerate bacterial numbers. Invasion rates were calculated as bacterial numbers recovered after the lysostaphin pulse (1 hour 10 minutes post-infection) relative to the initial bacterial inoculum.

Samples subjected to intracellular replication assessment were incubated with 20 µg/ml lysostaphin for 30 minutes, followed by washing and further incubation in medium containing 2 µg/ml lysostaphin until the end of the experiment. Intracellular replication was determined by plating dilution series as described above, at 3, 6 and 8 after infection.

### Live-cell imaging

HeLa or HeLa YFP-CWT cells were seeded in 8 well chamber µ-slides (ibidi) 24 hours prior to infection. Infection with fluorescent protein-expressing bacterial strains was performed at a MOI of 5 as described above. Time-lapse imaging of the samples was performed in imaging medium (RPMI1640 without phenol red containing 10% FBS and 2 µg/ml lysostaphin) on a Leica TCS SP5 confocal microscope using a 40x (Leica HC PL APO, NA=1.3) or 63x (Leica HCX PL APO, NA=1.3-0.6) oil immersion objective. The µ-slides were transferred to a pre-warmed live-cell incubation chamber (Life Imaging Systems) surrounding the confocal microscope and perfusion with 5 % CO_2_ in a humidified atmosphere and a temperature of 37 °C was applied during imaging. LAS AF software (Leica) was used for setting adjustment and image acquisition. All images were acquired at a resolution of 1024×1024 pixels and recorded in 8-bit mode at predefined time intervals. Z-stacks were imaged with a step size of 0.4 µm. All image-processing steps were performed using Fiji [97]. For detection of effector caspase activity 20 µM CellEvent Caspase 3/7 Green Detection Reagent (ThermoFisher Scientific) was added to the infected cells prior to imaging. Phase contrast microscopic images of live, infected cells were acquired with a LEICA DMR microscope connected to a SPOT camera using a 10x objective (Leica HC PL FLUOTAR, NA=0.32) and VisiView^®^ software (Visitron).

### Murine pneumonia infection

Overnight cultures of *S. aureus* strains in BHI medium were diluted to a final OD_600nm_ of 0.05 in 50 mL fresh BHI medium and grown for 3.5 hours at 37°C. All growth media for *S. aureus* strains containing plasmids were supplemented with appropriate antibiotics. After centrifugation, the bacterial pellet was resuspended in BHI with 20 % glycerol, aliquoted and stored at −80°C. For infection, aliquots were thawed, washed twice with PBS and adjusted to the desired infection inoculum of 2×10e^8^ CFU/20 µL. A sample was plated on TSB agar plates to confirm the correct bacterial concentration. Female Balb/c mice (6 weeks, Janvier Labs, Le Genest-Saint-Isle, France) were intranasally instilled with the infection dose. Mice were scored twice a day and sacrificed after 48 hours of infection. Lungs were harvested, homogenized and plated in serial dilutions on TSB agar plates in order to measure the bacterial burden in the individual organs.

### SDS-PAGE and Immunoblotting

For SDS-PAGE bacterial secreted proteins from sterile supernatant were precipitated overnight at −20 °C in 25 % trichloroacetic acid (TCA). After centrifugation at 4 °C for 30min the pellet was washed twice with ice-cold acetone. The dried pellet was resuspended in 2x Laemmli buffer (100 mM Tris/HCl (pH 6.8), 20 % glycerol, 4 % SDS, 1.5 % β-mercaptoethanol, 0.004% bromophenol blue) and immediately incubated at 95°C for 10 min for protein denaturation. Proteins were separated via gel electrophoresis on 12 % polyacrylamide gel and transferred to a PVDF membrane (Sigma Aldrich) using a semi-dry blotting system. The PVDF membrane was incubated for 1 hour in blocking solution (5 % human serum in 1x TBS-T) and overnight at 4°C with the first antibody (diluted in blocking solution). The staphopain A antibody (ABIN967004, antibodies online) was diluted 1:500 in blocking solution. α-toxin was detected with the respective antibody (S7531, Sigma Aldrich) diluted 1:2000 in blocking solution. Primary antibodies were detected with a horseradish peroxidase (HRP)-conjugated secondary antibody (170-6515, Biorad, 1:3000 in 1x TBS-T with 5 % non-fat dry milk) using enhanced chemiluminescence (ECL) and an Intas imaging system (Intas Science Imaging).

### Statistical analyses

Data were analyzed using GraphPad Prism Software (GraphPad Software, Version 6.01). For statistical analysis three biological replicates were performed, if not indicated otherwise. All data are presented as means with standard deviation (SD). P-values ≤0.05 were considered significant. Pairwise comparisons were assessed using unpaired Student’s t-test. Analysis of variance (ANOVA) was performed to determine whether a group of means was significantly different from each other. ANOVA was performed with Tukey’s post-hoc analysis for defining individual differences and Dunn’s multiple comparison test was applied for Kruskal-Wallis test.

## Acknowledgements

We thank the German Research Foundation (DFG; http://www.dfg.de) for funding this project within the Transregional Research Collaborative TRR34, project C11 (K.S., M.F., T.R.) and Z3 (T.H., K.O.). *S. aureus* JE2 mutants were obtained through the Network on Antimicrobial Resistance in *Staphylococcus aureus* (NARSA) Program supported under NIAID/ NIH Contract No. HHSN272200700055C. We are further grateful to Sudip Das for cloning of constructs, and Ursula Eilers and the Core Unit Functional Genomics (University Würzburg) for support with Operetta Imaging. Maria Steinke (University Hospital Würzburg and Fraunhofer IGB – Unit Würzburg) is thanked for technical advice during the establishment of the hTEC infection model. We thank Jan-Peter Hildebrandt (University Greifswald) for providing 16HBE14o^−^ cells.

## Ethics statement

All animal studies were approved by the local government of Franconia, Germany (approval number 55.2 2532-2-155) and performed in strict accordance with the guidelines for animal care and experimentation of German Animal Protection Law and the DIRECTIVE 2010/63/EU of the EU. The mice were housed in individually ventilated cages under normal diet in groups of four to five throughout the experiment with ad libitum access to food and water.

Bronchial segments for hTEC isolation were obtained from three patients undergoing elective pulmonary resection. Informed consent was obtained beforehand and the study was approved by the institutional ethics committee on human research of the Julius-Maximilians-University Würzburg (vote 182/10 and 17917) and Otto-von-Guericke University Magdeburg (vote 163/17).

## Author Contributions

KS, TH, AS, JP, KO, MJF and TR conceived and designed the experiments. KS, TH, AS, DK, HM, KP and AM performed the experiments. KS, TH, AM and KP analyzed the data. KS, MJF and TR wrote the paper.

## Supporting information

**S1 Table.**
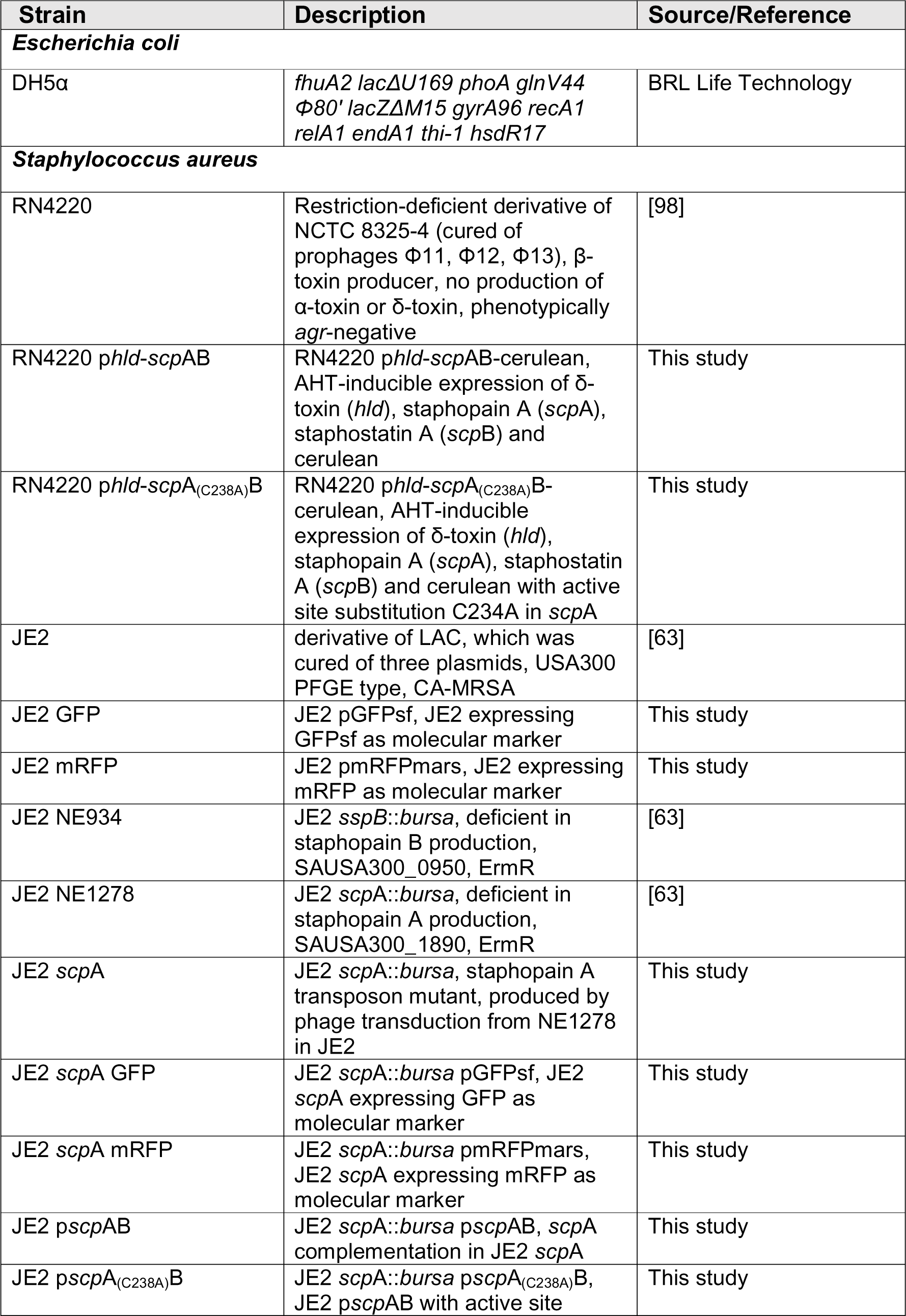
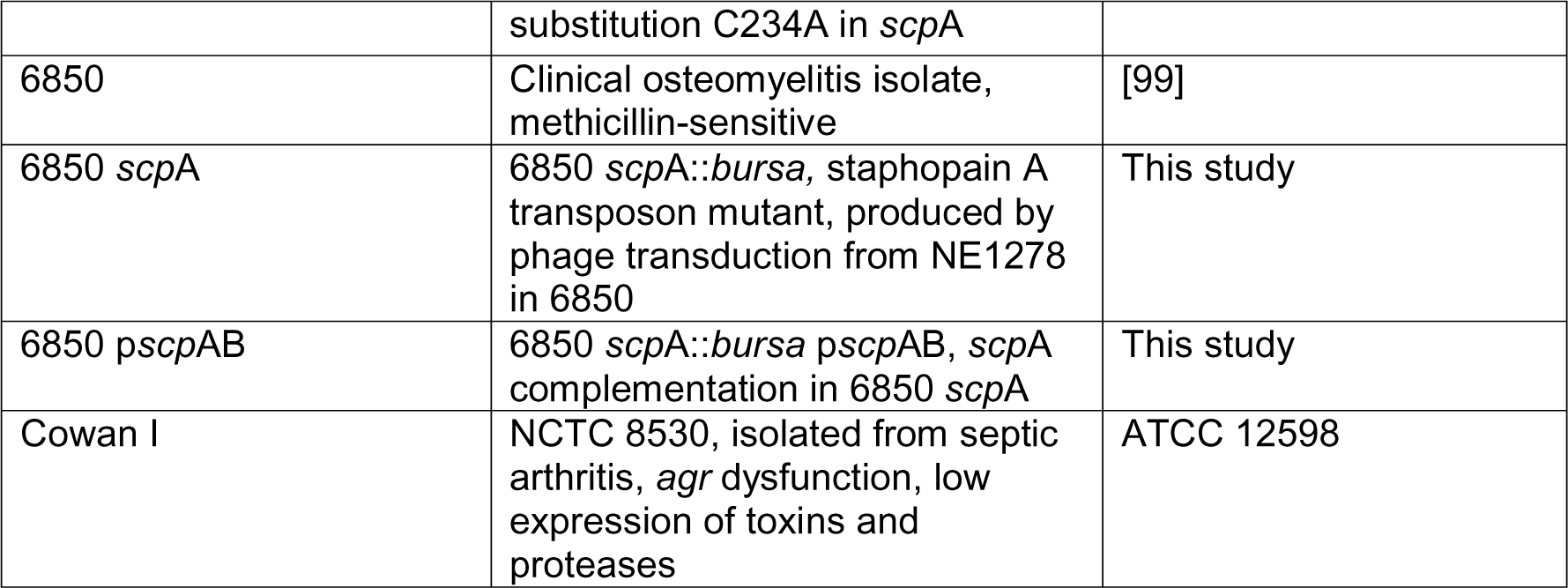
Bacterial strains used in this study.

**S2 Table.** Plasmids used in this study.

| Plasmid | Description | Reference |
| --- | --- | --- |
| p2085 | derivate of pALC2084 [100] with modified multiple-cloning site, shuttle vector, containing *tetR* and Pxyl/tet promoter driving GFPuvr expression, CmR | [101] |
| *pscpAB* | p2085-*scpAB*, complementation of *scpA*, native promotor of *scpAB* | This study |
| *pscpA*<sub>(C238A)</sub>B | *pscpAB* with active site substitution C234A in *scpA* | This study |
| *phld-hlb*-cerulean | p2085-*hld-hlb*-cerulean, AHT-inducible expression | [67] |
| *phld-scpAB*-cerulean | p2085-*hld-scpAB*-cerulean, AHT-inducible expression | This study |
| *phld-scpA*<sub>(C238A)</sub>B-cerulean | *phld-scpAB*-cerulean with active site substitution C234A in *scpA* | This study |
| pmRFPmars | p2085-SarAP1-mRFPmars, vector expressing mRFP in *S. aureus* under control of the constitutive sarAP1 promoter, CmR | [102] |
| pSK5632 | *S. aureus*-*E. coli* shuttle vector, low copy plasmid, CmR | [103] |
| pGFPsf | pSK5632-SarAP1-GFPsf, vector expressing GFPsf in *S. aureus* under control of the constitutive sarAP1 | This study |

**S3 Table 3.**
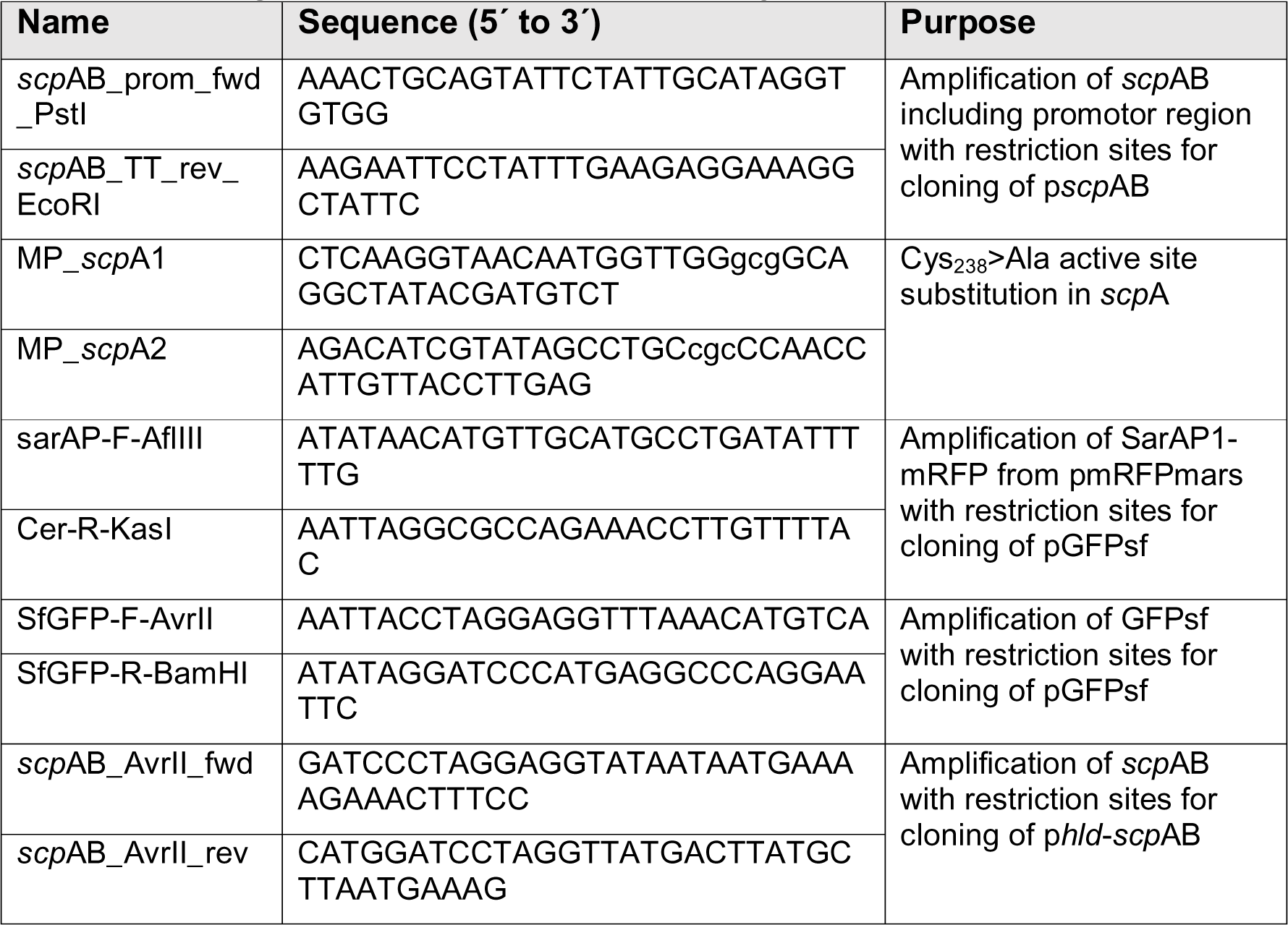
Oligonucleotides used in this study.

**S1 Fig. Staphopain A contributes to bacterial induced host cell death in *S. aureus* JE2 and 6850.** (A) HeLa cells were infected with *S. aureus* 6850 wild type, transposon mutant of staphopain A (6850 *scp*A) or complemented mutant (6850 p*scp*AB) and cytotoxicity was determined by LDH release 6 h p.i.. (B) HeLa cells were infected with wild type strain (JE2), staphopain A mutant (JE2 *scp*A), complemented mutant (JE2 p*scp*AB), complemented mutant with active site mutation (JE2 *scp*A_(C238A)_B) or Cowan I and cell death was assessed at 1.5, 3, 6, 12 and 24 h p.i. by LDH assay. (C, D) HeLa cells were infected with wild type (JE2), staphopain A mutant (JE2 *scp*A) or Cowan I and apoptotic cells were determined by annexin V-APC and 7AAD staining and flow cytometric analysis 6 h p.i.. Statistical significance was determined by one-way ANOVA (A) or two-way ANOVA (C) (*P<0.05, **P<0.01, ****P<0.0001).

**S2 Fig. Different morphologies of *S. aureus* JE2 and JE2 *scp*A-infected epithelial cells.** Microscopic images of A549, 16HBE14o^−^ and hTEC cells infected with JE2 wild type, JE2 *scp*A or Cowan I expressing mRFP at 6 (A549, 16HBE14o^−^) or 8 h p.i. (hTEC) (gray: PC, red: *S. aureus*, scale bar: 20 µm).

**S3 Fig. Invasion, phagosomal escape and intracellular replication of *S. aureus* JE2 and JE2 *scp*A in HeLa and 16HBE14o^−^ cells.** (A) HeLa cells were infected with JE2, JE2 *scp*A and Cowan I and invasion into HeLa cells was determined by quantifying intracellular CFUs at 1 h p.i.. (B) Phagosomal escape was quantified in the marker cell line 16HBE14o^−^ YFP-CWT 3 h p.i. by automated microscopy after infection with mRFP-expressing bacteria. (C) Infected HeLa YFP-CWT cells were imaged over time to visualize phagosomal escape of JE2 wild type (upper panel) and staphopain A mutant (JE2 *scp*A, lower panel) (red: *S. aureus*, yellow: YFP-CWT, gray: BF, scale bar: 20 µm). (D) Live cell imaging of 16HBE14o^−^ YFP-CWT cell infected with *S. aureus* JE2 mRFP or JE2 *scp*A mRFP (scale: 20 µm). (E) HeLa cells were infected with JE2, JE2 *scp*A and Cowan I and intracellular bacterial CFUs were quantified at 1, 3, 6 and 8 h p.i. Statistical significance was determined by unpaired t-test (B), one-way ANOVA (A) or two-way ANOVA (E) (**P<0.01).

**S4 Fig. Ectopic expression of staphopain A in a non-cytotoxic *S. aureus* strain leads to host cell death.** (A) Phase contrast images of infected HeLa cells at 2 h p.i. to reveal different morphologies of cells infected with RN4220 p*hld*-*scp*AB compared RN4220 p*hld*-*scp*A_(C238A)_B (scale bar: 100 µm). (B) The effect of E-64d (80 µM) pre-treatment on cytotoxicity of RN4220 p*hld*-*scp*AB infected HeLa cells was determined by quantification of LDH release 6 h p.i. and compared to solvent control (DMSO). (C) HeLa cells were infected with *S. aureus* RN4220 p*hld*-*scp*AB or RN4220 p*hld*-*scp*A_(C238A)_B with or without addition of 200 ng/ml AHT prior to infection. 4.5 h p.i. cells were stained with annexin V-APC and 7AAD and analyzed by flow cytometry. Statistical significance was determined by unpaired t test (B) or two-way ANOVA (C) (*P<0.05, **P<0.01, ***P<0.001, ****P<0.0001).

**S5 Fig. Maturation and enzymatic activity of staphopain A in sterile culture supernatant or of the purified protease.** (A) *S. aureus* RN4220 p*hld*-*scp*AB or RN4220 p*hld*-*scp*A_(C238A)_B were grown overnight with or without addition of 200 ng/ml AHT. Proteins of the sterile culture supernatant were precipitated and western blot was performed to detect staphopain A. Expression of functional ScpA was detected as the mature protein (triplet ranging from 17 to 20 kDa), while expression of a non-functional staphopain A led to accumulation of proScpA (ca. 40 kDa) [49]. (B, C) Proteolytic activity of staphopain A was measured from sterile culture supernatant of *S. aureus* RN4220 p*hld*-*scp*AB or RN4220 p*hld*-*scp*A_(C238A)_B (B) or increasing concentrations of the purified enzyme (C). Statistical significance was determined by unpaired t test (**P<0.01).

**S6 Fig. Cytotoxicity of *S. aureus* RN4220 p*hld*-*scp*AB and effect of protease inhibitors on staphopain A proteolytic activity.** (A) HeLa cells were infected with *S. aureus* RN4220 p*hld*-*scp*AB in the presence of 200 ng/ml AHT and 1 and 4.5 h p.i. cells were stained with annexin V-APC and 7AAD and analyzed by flow cytometry. (B) The inhibitors E-64 and Z-VAD-fmk reduce staphopain A proteolytic activity in a concentration-dependent manner. DMSO was used as solvent control. Statistical significance was determined by two-way ANOVA (****P<0.0001).

**S7 Fig. Growth curves of *S. aureus* JE2, JE2 *scp*A and JE2 p*scp*AB and** α**–toxin expression of *S. aureus* JE2 and JE2 *scp*A.** (A) *S. aureus* JE2, JE2 *scp*A and JE2 p*scp*AB were grown in TSB in 48-well plates for 18 h and the optical density at 600 nm was measured every 10 minutes. (B) *S. aureus* JE2 and JE2 *scp*A were grown overnight, proteins of the sterile culture supernatant were precipitated and western blot was performed to detect α-toxin (hla).

**S1 movie. Phagosomal escape of *S. aureus* JE2 and JE2 *scp*A.** HeLa YFP-CWT cells were infected with JE2 wild type (left panel) or staphopain a mutant (JE2 *scp*A, right panel) and imaged over time to visualize phagosomal escape (red: *S. aureus*, yellow: YFP-CWT, gray: BF).

**S2 movie. Intracellular replication of *S. aureus* JE2 and JE2 *scp*A.** HeLa cells were infected with *S. aureus* JE2 GFP (left panel) or JE2 *scp*A GFP (right panel) and time-lapse imaging was performed (green: *S. aureus*, gray: BF).

**S3 movie. Phagosomal escape of *S. aureus* RN4220 p*hld*-*scp*AB or RN4220 p*hld*-*scp*A_(C238A)_B.** HeLa YFP-CWT cells were infected with *S. aureus* p*hld*-*scp*AB (left panel) or RN4220 p*hld*-*scp*A_(C238A)_B (right panel) and time-lapse imaging was performed (cyan: *S. aureus*, yellow: YFP-CWT).

**S4 movie. Activation of effector caspases in HeLa cells infected with *S. aureus* RN4220 p*hld*-*scp*AB.** HeLa cells were infected with *S. aureus* RN4220 p*hld*-*scp*AB and a fluorogenic caspase3/7-substrate was added. Live cell imaging was performed to monitor the effect of staphopain A expression on cell morphology and activation of effectors caspases (cyan: *S. aureus*, green: CellEvent^TM^ Caspase3/7 Green Detection Reagent, gray: BF).

## References

1. Kluytmans J, van Belkum A, Verbrugh H. Nasal carriage of Staphylococcus aureus: epidemiology, underlying mechanisms, and associated risks. Clin Microbiol Rev. 1997;10(3):505–20. Epub 1997/07/01. PubMed PMID: 9227864; PubMed Central PMCID: PMCPMC172932.

2. Lowy FD. Staphylococcus aureus infections. N Engl J Med. 1998;339(8):520–32. Epub 1998/08/26. doi: 10.1056/NEJM199808203390806. PubMed PMID: 9709046.

3. Archer GL. Staphylococcus aureus: a well-armed pathogen. Clin Infect Dis. 1998;26(5):1179–81. Epub 1998/05/23. doi: 10.1086/520289. PubMed PMID: 9597249.

4. Nordmann P, Naas T, Fortineau N, Poirel L. Superbugs in the coming new decade; multidrug resistance and prospects for treatment of Staphylococcus aureus, Enterococcus spp. and Pseudomonas aeruginosa in 2010. Curr Opin Microbiol. 2007;10(5):436–40. Epub 2007/09/04. doi: 10.1016/j.mib.2007.07.004. PubMed PMID: 17765004.

5. Hamill RJ, Vann JM, Proctor RA. Phagocytosis of Staphylococcus aureus by cultured bovine aortic endothelial cells: model for postadherence events in endovascular infections. Infect Immun. 1986;54(3):833–6. Epub 1986/12/01. PubMed PMID: 3781627; PubMed Central PMCID: PMCPMC260245.

6. Jevon M, Guo C, Ma B, Mordan N, Nair SP, Harris M, et al. Mechanisms of internalization of Staphylococcus aureus by cultured human osteoblasts. Infect Immun. 1999;67(5):2677–81. Epub 1999/05/04. PubMed PMID: 10225942; PubMed Central PMCID: PMCPMC116025.

7. Kintarak S, Whawell SA, Speight PM, Packer S, Nair SP. Internalization of Staphylococcus aureus by human keratinocytes. Infect Immun. 2004;72(10):5668–75. Epub 2004/09/24. doi: 10.1128/IAI.72.10.5668-5675.2004. PubMed PMID: 15385465; PubMed Central PMCID: PMCPMC517534.

8. Strobel M, Pfortner H, Tuchscherr L, Volker U, Schmidt F, Kramko N, et al. Post-invasion events after infection with Staphylococcus aureus are strongly dependent on both the host cell type and the infecting S. aureus strain. Clin Microbiol Infect. 2016;22(9):799–809. Epub 2016/07/10. doi: 10.1016/j.cmi.2016.06.020. PubMed PMID: 27393124.

9. Kubica M, Guzik K, Koziel J, Zarebski M, Richter W, Gajkowska B, et al. A Potential New Pathway for Staphylococcus aureus Dissemination: The Silent Survival of S. aureus Phagocytosed by Human Monocyte-Derived Macrophages. Plos One. 2008;3(1). doi: 10.1371/journal.pone.0001409. PubMed PMID: WOS:000260469400006.

10. Blaettner S, Das S, Paprotka K, Eilers U, Krischke M, Kretschmer D, et al. Staphylococcus aureus Exploits a Non-ribosomal Cyclic Dipeptide to Modulate Survival within Epithelial Cells and Phagocytes. PLoS Pathog. 2016;12(9):e1005857. Epub 2016/09/16. doi: 10.1371/journal.ppat.1005857. PubMed PMID: 27632173; PubMed Central PMCID: PMCPMC5025175.

11. Flannagan RS, Heit B, Heinrichs DE. Intracellular replication of Staphylococcus aureus in mature phagolysosomes in macrophages precedes host cell death, and bacterial escape and dissemination. Cell Microbiol. 2016;18(4):514–35. Epub 2015/09/27. doi: 10.1111/cmi.12527. PubMed PMID: 26408990.

12. Gresham HD, Lowrance JH, Caver TE, Wilson BS, Cheung AL, Lindberg FP. Survival of Staphylococcus aureus inside neutrophils contributes to infection. J Immunol. 2000;164(7):3713–22. Epub 2000/03/22. PubMed PMID: 10725730.

13. Clement S, Vaudaux P, Francois P, Schrenzel J, Huggler E, Kampf S, et al. Evidence of an intracellular reservoir in the nasal mucosa of patients with recurrent Staphylococcus aureus rhinosinusitis. J Infect Dis. 2005;192(6):1023–8. Epub 2005/08/19. doi: 10.1086/432735. PubMed PMID: 16107955.

14. Hayes SM, Howlin R, Johnston DA, Webb JS, Clarke SC, Stoodley P, et al. Intracellular residency of Staphylococcus aureus within mast cells in nasal polyps: A novel observation. J Allergy Clin Immunol. 2015;135(6):1648–51. Epub 2015/02/15. doi: 10.1016/j.jaci.2014.12.1929. PubMed PMID: 25680455.

15. Hanssen AM, Kindlund B, Stenklev NC, Furberg AS, Fismen S, Olsen RS, et al. Localization of Staphylococcus aureus in tissue from the nasal vestibule in healthy carriers. BMC Microbiol. 2017;17(1):89. Epub 2017/04/07. doi: 10.1186/s12866-017-0997-3. PubMed PMID: 28381253; PubMed Central PMCID: PMCPMC5382455.

16. Li C, Wu Y, Riehle A, Ma J, Kamler M, Gulbins E, et al. Staphylococcus aureus Survives in Cystic Fibrosis Macrophages, Forming a Reservoir for Chronic Pneumonia. Infect Immun. 2017;85(5). Epub 2017/03/16. doi: 10.1128/IAI.00883-16. PubMed PMID: 28289144; PubMed Central PMCID: PMCPMC5400852.

17. Sinha B, Fraunholz M. Staphylococcus aureus host cell invasion and post-invasion events. Int J Med Microbiol. 2010;300(2-3):170–5. Epub 2009/09/29. doi: 10.1016/j.ijmm.2009.08.019. PubMed PMID: 19781990.

18. Horn J, Stelzner K, Rudel T, Fraunholz M. Inside job: Staphylococcus aureus host-pathogen interactions. Int J Med Microbiol. 2018;308(6):607–24. Epub 2017/12/09. doi: 10.1016/j.ijmm.2017.11.009. PubMed PMID: 29217333.

19. Moldovan A, Fraunholz MJ. In or out: phagosomal escape of Staphylococcus aureus. Cell Microbiol. 2018:e12997. Epub 2018/12/24. doi: 10.1111/cmi.12997. PubMed PMID: 30576050.

20. Jubrail J, Morris P, Bewley MA, Stoneham S, Johnston SA, Foster SJ, et al. Inability to sustain intraphagolysosomal killing of Staphylococcus aureus predisposes to bacterial persistence in macrophages. Cell Microbiol. 2016;18(1):80–96. Epub 2015/08/08. doi: 10.1111/cmi.12485. PubMed PMID: 26248337; PubMed Central PMCID: PMCPMC4778410.

21. Thwaites GE, Gant V. Are bloodstream leukocytes Trojan Horses for the metastasis of Staphylococcus aureus? Nat Rev Microbiol. 2011;9(3):215–22. Epub 2011/02/08. doi: 10.1038/nrmicro2508. PubMed PMID: 21297670.

22. Lehar SM, Pillow T, Xu M, Staben L, Kajihara KK, Vandlen R, et al. Novel antibody-antibiotic conjugate eliminates intracellular S. aureus. Nature. 2015;527(7578):323-8. Epub 2015/11/05. doi: 10.1038/nature16057. PubMed PMID: 26536114.

23. Flieger A, Frischknecht F, Hacker G, Hornef MW, Pradel G. Pathways of host cell exit by intracellular pathogens. Microb Cell. 2018;5(12):525–44. Epub 2018/12/12. doi: 10.15698/mic2018.12.659. PubMed PMID: 30533418; PubMed Central PMCID: PMCPMC6282021.

24. Young AB, Cooley ID, Chauhan VS, Marriott I. Causative agents of osteomyelitis induce death domain-containing TNF-related apoptosis-inducing ligand receptor expression on osteoblasts. Bone. 2011;48(4):857–63. Epub 2010/12/07. doi: 10.1016/j.bone.2010.11.015. PubMed PMID: 21130908.

25. Kahl BC, Goulian M, van Wamel W, Herrmann M, Simon SM, Kaplan G, et al. Staphylococcus aureus RN6390 replicates and induces apoptosis in a pulmonary epithelial cell line. Infect Immun. 2000;68(9):5385–92. Epub 2000/08/19. PubMed PMID: 10948168; PubMed Central PMCID: PMCPMC101802.

26. Weglarczyk K, Baran J, Zembala M, Pryjma J. Caspase-8 activation precedes alterations of mitochondrial membrane potential during monocyte apoptosis induced by phagocytosis and killing of Staphylococcus aureus. Infect Immun. 2004;72(5):2590–7. Epub 2004/04/23. PubMed PMID: 15102767; PubMed Central PMCID: PMCPMC387870.

27. Nuzzo I, Sanges MR, Folgore A, Carratelli CR. Apoptosis of human keratinocytes after bacterial invasion. FEMS Immunol Med Microbiol. 2000;27(3):235–40. Epub 2000/02/23. doi: 10.1111/j.1574-695X.2000.tb01435.x. PubMed PMID: 10683468.

28. Haslinger-Loeffler B, Wagner B, Bruck M, Strangfeld K, Grundmeier M, Fischer U, et al. Staphylococcus aureus induces caspase-independent cell death in human peritoneal mesothelial cells. Kidney Int. 2006;70(6):1089–98. Epub 2006/07/28. doi: 10.1038/sj.ki.5001710. PubMed PMID: 16871245.

29. Schnaith A, Kashkar H, Leggio SA, Addicks K, Kronke M, Krut O. Staphylococcus aureus subvert autophagy for induction of caspase-independent host cell death. J Biol Chem. 2007;282(4):2695–706. Epub 2006/12/01. doi: 10.1074/jbc.M609784200. PubMed PMID: 17135247.

30. Kobayashi SD, Braughton KR, Palazzolo-Ballance AM, Kennedy AD, Sampaio E, Kristosturyan E, et al. Rapid neutrophil destruction following phagocytosis of Staphylococcus aureus. J Innate Immun. 2010;2(6):560–75. Epub 2010/07/01. doi: 10.1159/000317134. PubMed PMID: 20587998; PubMed Central PMCID: PMCPMC3219502.

31. Greenlee-Wacker MC, Rigby KM, Kobayashi SD, Porter AR, DeLeo FR, Nauseef WM. Phagocytosis of Staphylococcus aureus by human neutrophils prevents macrophage efferocytosis and induces programmed necrosis. J Immunol. 2014;192(10):4709–17. Epub 2014/04/15. doi: 10.4049/jimmunol.1302692. PubMed PMID: 24729616; PubMed Central PMCID: PMCPMC4011196.

32. Menzies BE, Kourteva I. Staphylococcus aureus alpha-toxin induces apoptosis in endothelial cells. FEMS Immunol Med Microbiol. 2000;29(1):39–45. Epub 2000/09/01. doi: 10.1111/j.1574-695X.2000.tb01503.x. PubMed PMID: 10967259.

33. Haslinger-Loeffler B, Kahl BC, Grundmeier M, Strangfeld K, Wagner B, Fischer U, et al. Multiple virulence factors are required for Staphylococcus aureus-induced apoptosis in endothelial cells. Cell Microbiol. 2005;7(8):1087–97. Epub 2005/07/13. doi: 10.1111/j.1462-5822.2005.00533.x. PubMed PMID: 16008576.

34. Melehani JH, James DB, DuMont AL, Torres VJ, Duncan JA. Staphylococcus aureus Leukocidin A/B (LukAB) Kills Human Monocytes via Host NLRP3 and ASC when Extracellular, but Not Intracellular. PLoS Pathog. 2015;11(6):e1004970. Epub 2015/06/13. doi: 10.1371/journal.ppat.1004970. PubMed PMID: 26069969; PubMed Central PMCID: PMCPMC4466499.

35. Muenzenmayer L, Geiger T, Daiber E, Schulte B, Autenrieth SE, Fraunholz M, et al. Influence of Sae-regulated and Agr-regulated factors on the escape of Staphylococcus aureus from human macrophages. Cell Microbiol. 2016;18(8):1172–83. Epub 2016/02/21. doi: 10.1111/cmi.12577. PubMed PMID: 26895738.

36. Ventura CL, Malachowa N, Hammer CH, Nardone GA, Robinson MA, Kobayashi SD, et al. Identification of a novel Staphylococcus aureus two-component leukotoxin using cell surface proteomics. Plos One. 2010;5(7):e11634. Epub 2010/07/28. doi: 10.1371/journal.pone.0011634. PubMed PMID: 20661294; PubMed Central PMCID: PMCPMC2905442.

37. DuMont AL, Yoong P, Surewaard BG, Benson MA, Nijland R, van Strijp JA, et al. Staphylococcus aureus elaborates leukocidin AB to mediate escape from within human neutrophils. Infect Immun. 2013;81(5):1830–41. Epub 2013/03/20. doi: 10.1128/IAI.00095-13. PubMed PMID: 23509138; PubMed Central PMCID: PMCPMC3648020.

38. Chi CY, Lin CC, Liao IC, Yao YC, Shen FC, Liu CC, et al. Panton-Valentine leukocidin facilitates the escape of Staphylococcus aureus from human keratinocyte endosomes and induces apoptosis. J Infect Dis. 2014;209(2):224–35. Epub 2013/08/21. doi: 10.1093/infdis/jit445. PubMed PMID: 23956440.

39. Jin T, Zhu YL, Li J, Shi J, He XQ, Ding J, et al. Staphylococcal protein A, Panton-Valentine leukocidin and coagulase aggravate the bone loss and bone destruction in osteomyelitis. Cell Physiol Biochem. 2013;32(2):322–33. Epub 2013/08/15. doi: 10.1159/000354440. PubMed PMID: 23942321.

40. Rasigade JP, Trouillet-Assant S, Ferry T, Diep BA, Sapin A, Lhoste Y, et al. PSMs of hypervirulent Staphylococcus aureus act as intracellular toxins that kill infected osteoblasts. Plos One. 2013;8(5):e63176. Epub 2013/05/22. doi: 10.1371/journal.pone.0063176. PubMed PMID: 23690994; PubMed Central PMCID: PMCPMC3653922.

41. Surewaard BG, de Haas CJ, Vervoort F, Rigby KM, DeLeo FR, Otto M, et al. Staphylococcal alpha-phenol soluble modulins contribute to neutrophil lysis after phagocytosis. Cell Microbiol. 2013;15(8):1427–37. Epub 2013/03/09. doi: 10.1111/cmi.12130. PubMed PMID: 23470014; PubMed Central PMCID: PMCPMC4784422.

42. Dubin G. Extracellular proteases of Staphylococcus spp. Biol Chem. 2002;383(7-8):1075–86. Epub 2002/11/20. doi: 10.1515/BC.2002.116. PubMed PMID: 12437090.

43. Hofmann B, Schomburg D, Hecht HJ. Crystal structure of a thiol proteinase from Staphylococcus aureus V-8 in the E-64 inhibitor complex. Acta Crystallographica. 1993;49 (Supplement):102.

44. Arvidson S. Extracellular enzymes. In: Fischetti VA, Novick RP, Ferretti JJ, Potrnoy DA, Rood JI, editors. Gram-positive pathogens. Washington, D.C.: American Society for Microbiology; 2000. p. 379–85.

45. Golonka E, Filipek R, Sabat A, Sinczak A, Potempa J. Genetic characterization of staphopain genes in Staphylococcus aureus. Biol Chem. 2004;385(11):1059–67. Epub 2004/12/04. doi: 10.1515/BC.2004.137. PubMed PMID: 15576326.

46. Rzychon M, Sabat A, Kosowska K, Potempa J, Dubin A. Staphostatins: an expanding new group of proteinase inhibitors with a unique specificity for the regulation of staphopains, Staphylococcus spp. cysteine proteinases. Mol Microbiol. 2003;49(4):1051–66. Epub 2003/08/02. PubMed PMID: 12890028.

47. Massimi I, Park E, Rice K, Muller-Esterl W, Sauder D, McGavin MJ. Identification of a novel maturation mechanism and restricted substrate specificity for the SspB cysteine protease of Staphylococcus aureus. J Biol Chem. 2002;277(44):41770–7. Epub 2002/09/19. doi: 10.1074/jbc.M207162200. PubMed PMID: 12207024.

48. Filipek R, Rzychon M, Oleksy A, Gruca M, Dubin A, Potempa J, et al. The Staphostatin-staphopain complex: a forward binding inhibitor in complex with its target cysteine protease. J Biol Chem. 2003;278(42):40959–66. Epub 2003/07/23. doi: 10.1074/jbc.M302926200. PubMed PMID: 12874290.

49. Nickerson N, Ip J, Passos DT, McGavin MJ. Comparison of Staphopain A (ScpA) and B (SspB) precursor activation mechanisms reveals unique secretion kinetics of proSspB (Staphopain B), and a different interaction with its cognate Staphostatin, SspC. Mol Microbiol. 2010;75(1):161–77. Epub 2009/12/01. doi: 10.1111/j.1365-2958.2009.06974.x. PubMed PMID: 19943908.

50. Drapeau GR. Role of metalloprotease in activation of the precursor of staphylococcal protease. J Bacteriol. 1978;136(2):607–13. Epub 1978/11/01. PubMed PMID: 711676; PubMed Central PMCID: PMCPMC218585.

51. Bjoerklind A, Joernvall H. Substrate specificity of three different extracellular proteolytic enzymes from Staphylococcus aureus. Biochim Biophys Acta. 1974;370(2):524–9. Epub 1974/12/29. doi: 10.1016/0005-2744(74)90113-2. PubMed PMID: 4613383.

52. Ohbayashi T, Irie A, Murakami Y, Nowak M, Potempa J, Nishimura Y, et al. Degradation of fibrinogen and collagen by staphopains, cysteine proteases released from Staphylococcus aureus. Microbiology. 2011;157(Pt 3):786–92. Epub 2010/11/18. doi: 10.1099/mic.0.044503-0. PubMed PMID: 21081759.

53. Laarman AJ, Mijnheer G, Mootz JM, van Rooijen WJ, Ruyken M, Malone CL, et al. Staphylococcus aureus Staphopain A inhibits CXCR2-dependent neutrophil activation and chemotaxis. EMBO J. 2012;31(17):3607–19. Epub 2012/08/02. doi: 10.1038/emboj.2012.212. PubMed PMID: 22850671; PubMed Central PMCID: PMCPMC3433787.

54. Smagur J, Guzik K, Bzowska M, Kuzak M, Zarebski M, Kantyka T, et al. Staphylococcal cysteine protease staphopain B (SspB) induces rapid engulfment of human neutrophils and monocytes by macrophages. Biol Chem. 2009;390(4):361–71. Epub 2009/03/17. doi: 10.1515/BC.2009.042. PubMed PMID: 19284294.

55. Imamura T, Tanase S, Szmyd G, Kozik A, Travis J, Potempa J. Induction of vascular leakage through release of bradykinin and a novel kinin by cysteine proteinases from Staphylococcus aureus. J Exp Med. 2005;201(10):1669–76. Epub 2005/05/18. doi: 10.1084/jem.20042041. PubMed PMID: 15897280; PubMed Central PMCID: PMCPMC2212919.

56. Mootz JM, Malone CL, Shaw LN, Horswill AR. Staphopains modulate Staphylococcus aureus biofilm integrity. Infect Immun. 2013;81(9):3227–38. Epub 2013/06/27. doi: 10.1128/IAI.00377-13. PubMed PMID: 23798534; PubMed Central PMCID: PMCPMC3754231.

57. Potempa J, Dubin A, Korzus G, Travis J. Degradation of elastin by a cysteine proteinase from Staphylococcus aureus. J Biol Chem. 1988;263(6):2664–7. Epub 1988/02/25. PubMed PMID: 3422637.

58. Sloot N, Thomas M, Marre R, Gatermann S. Purification and characterisation of elastase from Staphylococcus epidermidis. J Med Microbiol. 1992;37(3):201–5. Epub 1992/09/01. doi: 10.1099/00222615-37-3-201. PubMed PMID: 1518036.

59. Martinez-Garcia S, Rodriguez-Martinez S, Cancino-Diaz ME, Cancino-Diaz JC. Extracellular proteases of Staphylococcus epidermidis: roles as virulence factors and their participation in biofilm. APMIS. 2018;126(3):177–85. Epub 2018/02/06. doi: 10.1111/apm.12805. PubMed PMID: 29399876.

60. Yokoi K, Kakikawa M, Kimoto H, Watanabe K, Yasukawa H, Yamakawa A, et al. Genetic and biochemical characterization of glutamyl endopeptidase of Staphylococcus warneri M. Gene. 2001;281(1-2):115–22. Epub 2001/12/26. doi: 10.1016/s0378-1119(01)00782-x. PubMed PMID: 11750133.

61. Dubin G, Wladyka B, Stec-Niemczyk J, Chmiel D, Zdzalik M, Dubin A, et al. The staphostatin family of cysteine protease inhibitors in the genus Staphylococcus as an example of parallel evolution of protease and inhibitor specificity. Biol Chem. 2007;388(2):227–35. Epub 2007/01/31. doi: 10.1515/BC.2007.025. PubMed PMID: 17261086.

62. Burd JF, Usategui-Gomez M. A colorimetric assay for serum lactate dehydrogenase. Clin Chim Acta. 1973;46(3):223–7. Epub 1973/07/14. PubMed PMID: 4725386.

63. Fey PD, Endres JL, Yajjala VK, Widhelm TJ, Boissy RJ, Bose JL, et al. A genetic resource for rapid and comprehensive phenotype screening of nonessential Staphylococcus aureus genes. MBio. 2013;4(1):e00537–12. Epub 2013/02/14. doi: 10.1128/mBio.00537-12. PubMed PMID: 23404398; PubMed Central PMCID: PMCPMC3573662.

64. Kroemer G, Galluzzi L, Vandenabeele P, Abrams J, Alnemri ES, Baehrecke EH, et al. Classification of cell death: recommendations of the Nomenclature Committee on Cell Death 2009. Cell Death Differ. 2009;16(1):3–11. Epub 2008/10/11. doi: 10.1038/cdd.2008.150. PubMed PMID: 18846107; PubMed Central PMCID: PMCPMC2744427.

65. Vermes I, Haanen C, Steffens-Nakken H, Reutelingsperger C. A novel assay for apoptosis. Flow cytometric detection of phosphatidylserine expression on early apoptotic cells using fluorescein labelled Annexin V. J Immunol Methods. 1995;184(1):39–51. Epub 1995/07/17. PubMed PMID: 7622868.

66. Grosz M, Kolter J, Paprotka K, Winkler AC, Schafer D, Chatterjee SS, et al. Cytoplasmic replication of Staphylococcus aureus upon phagosomal escape triggered by phenol-soluble modulin alpha. Cell Microbiol. 2014;16(4):451–65. Epub 2013/10/30. doi: 10.1111/cmi.12233. PubMed PMID: 24164701; PubMed Central PMCID: PMCPMC3969633.

67. Giese B, Glowinski F, Paprotka K, Dittmann S, Steiner T, Sinha B, et al. Expression of delta-toxin by Staphylococcus aureus mediates escape from phago-endosomes of human epithelial and endothelial cells in the presence of beta-toxin. Cell Microbiol. 2011;13(2):316–29. Epub 2010/10/16. doi: 10.1111/j.1462-5822.2010.01538.x. PubMed PMID: 20946243.

68. Cohen GM. Caspases: the executioners of apoptosis. Biochem J. 1997;326 (Pt 1):1–16. Epub 1997/08/15. PubMed PMID: 9337844; PubMed Central PMCID: PMCPMC1218630.

69. Bubeck Wardenburg J, Bae T, Otto M, Deleo FR, Schneewind O. Poring over pores: alpha-hemolysin and Panton-Valentine leukocidin in Staphylococcus aureus pneumonia. Nat Med. 2007;13(12):1405–6. Epub 2007/12/08. doi: 10.1038/nm1207-1405. PubMed PMID: 18064027.

70. Thomas JA, Tan MSY, Bisson C, Borg A, Umrekar TR, Hackett F, et al. A protease cascade regulates release of the human malaria parasite Plasmodium falciparum from host red blood cells. Nat Microbiol. 2018;3(4):447–55. Epub 2018/02/21. doi: 10.1038/s41564-018-0111-0. PubMed PMID: 29459732; PubMed Central PMCID: PMCPMC6089347.

71. Huai J, Jockel L, Schrader K, Borner C. Role of caspases and non-caspase proteases in cell death. F1000 Biol Rep. 2010;2. Epub 2010/10/16. doi: 10.3410/B2-48. PubMed PMID: 20948786; PubMed Central PMCID: PMCPMC2950037.

72. Shaw L, Golonka E, Potempa J, Foster SJ. The role and regulation of the extracellular proteases of Staphylococcus aureus. Microbiology. 2004;150(Pt 1):217–28. Epub 2004/01/02. doi: 10.1099/mic.0.26634-0. PubMed PMID: 14702415.

73. Calander AM, Jonsson IM, Kanth A, Arvidsson S, Shaw L, Foster SJ, et al. Impact of staphylococcal protease expression on the outcome of infectious arthritis. Microbes Infect. 2004;6(2):202–6. Epub 2004/03/05. doi: 10.1016/j.micinf.2003.10.015. PubMed PMID: 14998519.

74. Zhang Z, Abdel-Razek O, Hawgood S, Wang G. Protective Role of Surfactant Protein D in Ocular Staphylococcus aureus Infection. Plos One. 2015;10(9):e0138597. Epub 2015/09/24. doi: 10.1371/journal.pone.0138597. PubMed PMID: 26398197; PubMed Central PMCID: PMCPMC4580580.

75. Frees D, Brondsted L, Ingmer H. Bacterial proteases and virulence. Subcell Biochem. 2013;66:161–92. Epub 2013/03/13. doi: 10.1007/978-94-007-5940-4_7. PubMed PMID: 23479441.

76. Lam TT, Giese B, Chikkaballi D, Kuhn A, Wolber W, Pane-Farre J, et al. Phagolysosomal integrity is generally maintained after Staphylococcus aureus invasion of nonprofessional phagocytes but is modulated by strain 6850. Infect Immun. 2010;78(8):3392–403. Epub 2010/06/10. doi: 10.1128/IAI.00012-10. PubMed PMID: 20530231; PubMed Central PMCID: PMCPMC2916288.

77. Das S, Lindemann C, Young BC, Muller J, Osterreich B, Ternette N, et al. Natural mutations in a Staphylococcus aureus virulence regulator attenuate cytotoxicity but permit bacteremia and abscess formation. Proc Natl Acad Sci U S A. 2016;113(22):E3101–10. Epub 2016/05/18. doi: 10.1073/pnas.1520255113. PubMed PMID: 27185949; PubMed Central PMCID: PMCPMC4896717.

78. Maeder U, Nicolas P, Depke M, Pane-Farre J, Debarbouille M, van der Kooi-Pol MM, et al. Staphylococcus aureus Transcriptome Architecture: From Laboratory to Infection-Mimicking Conditions. PLoS Genet. 2016;12(4):e1005962. Epub 2016/04/02. doi: 10.1371/journal.pgen.1005962. PubMed PMID: 27035918; PubMed Central PMCID: PMCPMC4818034.

79. Traber K, Novick R. A slipped-mispairing mutation in AgrA of laboratory strains and clinical isolates results in delayed activation of agr and failure to translate delta- and alpha-haemolysins. Mol Microbiol. 2006;59(5):1519–30. Epub 2006/02/14. doi: 10.1111/j.1365-2958.2006.04986.x. PubMed PMID: 16468992.

80. Nair D, Memmi G, Hernandez D, Bard J, Beaume M, Gill S, et al. Whole-genome sequencing of Staphylococcus aureus strain RN4220, a key laboratory strain used in virulence research, identifies mutations that affect not only virulence factors but also the fitness of the strain. J Bacteriol. 2011;193(9):2332–5. Epub 2011/03/08. doi: 10.1128/JB.00027-11. PubMed PMID: 21378186; PubMed Central PMCID: PMCPMC3133102.

81. Verdon J, Girardin N, Lacombe C, Berjeaud JM, Hechard Y. delta-hemolysin, an update on a membrane-interacting peptide. Peptides. 2009;30(4):817–23. Epub 2009/01/20. doi: 10.1016/j.peptides.2008.12.017. PubMed PMID: 19150639.

82. Smagur J, Guzik K, Magiera L, Bzowska M, Gruca M, Thogersen IB, et al. A new pathway of staphylococcal pathogenesis: apoptosis-like death induced by Staphopain B in human neutrophils and monocytes. J Innate Immun. 2009;1(2):98–108. Epub 2009/01/01. doi: 10.1159/000181014. PubMed PMID: 20375568.

83. Martin SJ, Reutelingsperger CP, McGahon AJ, Rader JA, van Schie RC, LaFace DM, et al. Early redistribution of plasma membrane phosphatidylserine is a general feature of apoptosis regardless of the initiating stimulus: inhibition by overexpression of Bcl-2 and Abl. J Exp Med. 1995;182(5):1545–56. Epub 1995/11/01. PubMed PMID: 7595224; PubMed Central PMCID: PMCPMC2192182.

84. Lynch C, Panagopoulou M, Gregory CD. Extracellular Vesicles Arising from Apoptotic Cells in Tumors: Roles in Cancer Pathogenesis and Potential Clinical Applications. Front Immunol. 2017;8:1174. Epub 2017/10/12. doi: 10.3389/fimmu.2017.01174. PubMed PMID: 29018443; PubMed Central PMCID: PMCPMC5614926.

85. Vanden Berghe T, Vanlangenakker N, Parthoens E, Deckers W, Devos M, Festjens N, et al. Necroptosis, necrosis and secondary necrosis converge on similar cellular disintegration features. Cell Death Differ. 2010;17(6):922–30. Epub 2009/12/17. doi: 10.1038/cdd.2009.184. PubMed PMID: 20010783.

86. Kalinska M, Kantyka T, Greenbaum DC, Larsen KS, Wladyka B, Jabaiah A, et al. Substrate specificity of Staphylococcus aureus cysteine proteases--Staphopains A, B and C. Biochimie. 2012;94(2):318–27. Epub 2011/08/02. doi: 10.1016/j.biochi.2011.07.020. PubMed PMID: 21802486.

87. Cirman T, Oresic K, Mazovec GD, Turk V, Reed JC, Myers RM, et al. Selective disruption of lysosomes in HeLa cells triggers apoptosis mediated by cleavage of Bid by multiple papain-like lysosomal cathepsins. J Biol Chem. 2004;279(5):3578–87. Epub 2003/10/29. doi: 10.1074/jbc.M308347200. PubMed PMID: 14581476.

88. Droga-Mazovec G, Bojic L, Petelin A, Ivanova S, Romih R, Repnik U, et al. Cysteine cathepsins trigger caspase-dependent cell death through cleavage of bid and antiapoptotic Bcl-2 homologues. J Biol Chem. 2008;283(27):19140–50. Epub 2008/05/13. doi: 10.1074/jbc.M802513200. PubMed PMID: 18469004.

89. Roberg K, Kagedal K, Ollinger K. Microinjection of cathepsin d induces caspase-dependent apoptosis in fibroblasts. Am J Pathol. 2002;161(1):89–96. Epub 2002/07/11. doi: 10.1016/S0002-9440(10)64160-0. PubMed PMID: 12107093; PubMed Central PMCID: PMCPMC1850710.

90. Stoka V, Turk B, Schendel SL, Kim TH, Cirman T, Snipas SJ, et al. Lysosomal protease pathways to apoptosis. Cleavage of bid, not pro-caspases, is the most likely route. J Biol Chem. 2001;276(5):3149–57. Epub 2000/11/14. doi: 10.1074/jbc.M008944200. PubMed PMID: 11073962.

91. Vincents B, Onnerfjord P, Gruca M, Potempa J, Abrahamson M. Down-regulation of human extracellular cysteine protease inhibitors by the secreted staphylococcal cysteine proteases, staphopain A and B. Biol Chem. 2007;388(4):437–46. Epub 2007/03/30. doi: 10.1515/BC.2007.042. PubMed PMID: 17391065.

92. Kolar SL, Ibarra JA, Rivera FE, Mootz JM, Davenport JE, Stevens SM, et al. Extracellular proteases are key mediators of Staphylococcus aureus virulence via the global modulation of virulence-determinant stability. Microbiologyopen. 2013;2(1):18–34. Epub 2012/12/13. doi: 10.1002/mbo3.55. PubMed PMID: 23233325; PubMed Central PMCID: PMCPMC3584211.

93. Szafranska AK, Oxley AP, Chaves-Moreno D, Horst SA, Rosslenbroich S, Peters G, et al. High-resolution transcriptomic analysis of the adaptive response of Staphylococcus aureus during acute and chronic phases of osteomyelitis. MBio. 2014;5(6). Epub 2014/12/30. doi: 10.1128/mBio.01775-14. PubMed PMID: 25538190; PubMed Central PMCID: PMCPMC4278534.

94. Kantyka T, Pyrc K, Gruca M, Smagur J, Plaza K, Guzik K, et al. Staphylococcus aureus proteases degrade lung surfactant protein A potentially impairing innate immunity of the lung. J Innate Immun. 2013;5(3):251–60. Epub 2012/12/14. doi: 10.1159/000345417. PubMed PMID: 23235402; PubMed Central PMCID: PMCPMC3787703.

95. Cozens AL, Yezzi MJ, Kunzelmann K, Ohrui T, Chin L, Eng K, et al. CFTR expression and chloride secretion in polarized immortal human bronchial epithelial cells. Am J Respir Cell Mol Biol. 1994;10(1):38–47. Epub 1994/01/01. doi: 10.1165/ajrcmb.10.1.7507342. PubMed PMID: 7507342.

96. Steinke M, Gross R, Walles H, Gangnus R, Schutze K, Walles T. An engineered 3D human airway mucosa model based on an SIS scaffold. Biomaterials. 2014;35(26):7355–62. Epub 2014/06/11. doi: 10.1016/j.biomaterials.2014.05.031. PubMed PMID: 24912816.

97. Schindelin J, Arganda-Carreras I, Frise E, Kaynig V, Longair M, Pietzsch T, et al. Fiji: an open-source platform for biological-image analysis. Nat Methods. 2012;9(7):676–82. Epub 2012/06/30. doi: 10.1038/nmeth.2019. PubMed PMID: 22743772; PubMed Central PMCID: PMCPMC3855844.

98. Kreiswirth BN, Lofdahl S, Betley MJ, O’Reilly M, Schlievert PM, Bergdoll MS, et al. The toxic shock syndrome exotoxin structural gene is not detectably transmitted by a prophage. Nature. 1983;305(5936):709–12. Epub 1983/10/20. PubMed PMID: 6226876.

99. Vann JM, Proctor RA. Ingestion of Staphylococcus aureus by bovine endothelial cells results in time- and inoculum-dependent damage to endothelial cell monolayers. Infect Immun. 1987;55(9):2155–63. Epub 1987/09/01. PubMed PMID: 3623696; PubMed Central PMCID: PMCPMC260672.

100. Bateman BT, Donegan NP, Jarry TM, Palma M, Cheung AL. Evaluation of a tetracycline-inducible promoter in Staphylococcus aureus in vitro and in vivo and its application in demonstrating the role of sigB in microcolony formation. Infect Immun. 2001;69(12):7851–7. Epub 2001/11/14. doi: 10.1128/IAI.69.12.7851-7857.2001. PubMed PMID: 11705967; PubMed Central PMCID: PMCPMC98881.

101. Giese B, Dittmann S, Paprotka K, Levin K, Weltrowski A, Biehler D, et al. Staphylococcal alpha-toxin is not sufficient to mediate escape from phagolysosomes in upper-airway epithelial cells. Infect Immun. 2009;77(9):3611–25. Epub 2009/07/01. doi: 10.1128/IAI.01478-08. PubMed PMID: 19564384; PubMed Central PMCID: PMCPMC2738027.

102. Paprotka K, Giese B, Fraunholz MJ. Codon-improved fluorescent proteins in investigation of Staphylococcus aureus host pathogen interactions. J Microbiol Methods. 2010;83(1):82–6. Epub 2010/08/17. doi: 10.1016/j.mimet.2010.07.022. PubMed PMID: 20708040.

103. Grkovic S, Brown MH, Hardie KM, Firth N, Skurray RA. Stable low-copy-number Staphylococcus aureus shuttle vectors. Microbiology. 2003;149(Pt 3):785–94. Epub 2003/03/14. doi: 10.1099/mic.0.25951-0. PubMed PMID: 12634346.

