## Supplementary figures and images for "Intracellular *Staphylococcus aureus* employs the cysteine protease staphopain A to induce host cell death in epithelial cells"

### Supplemental Figure 1

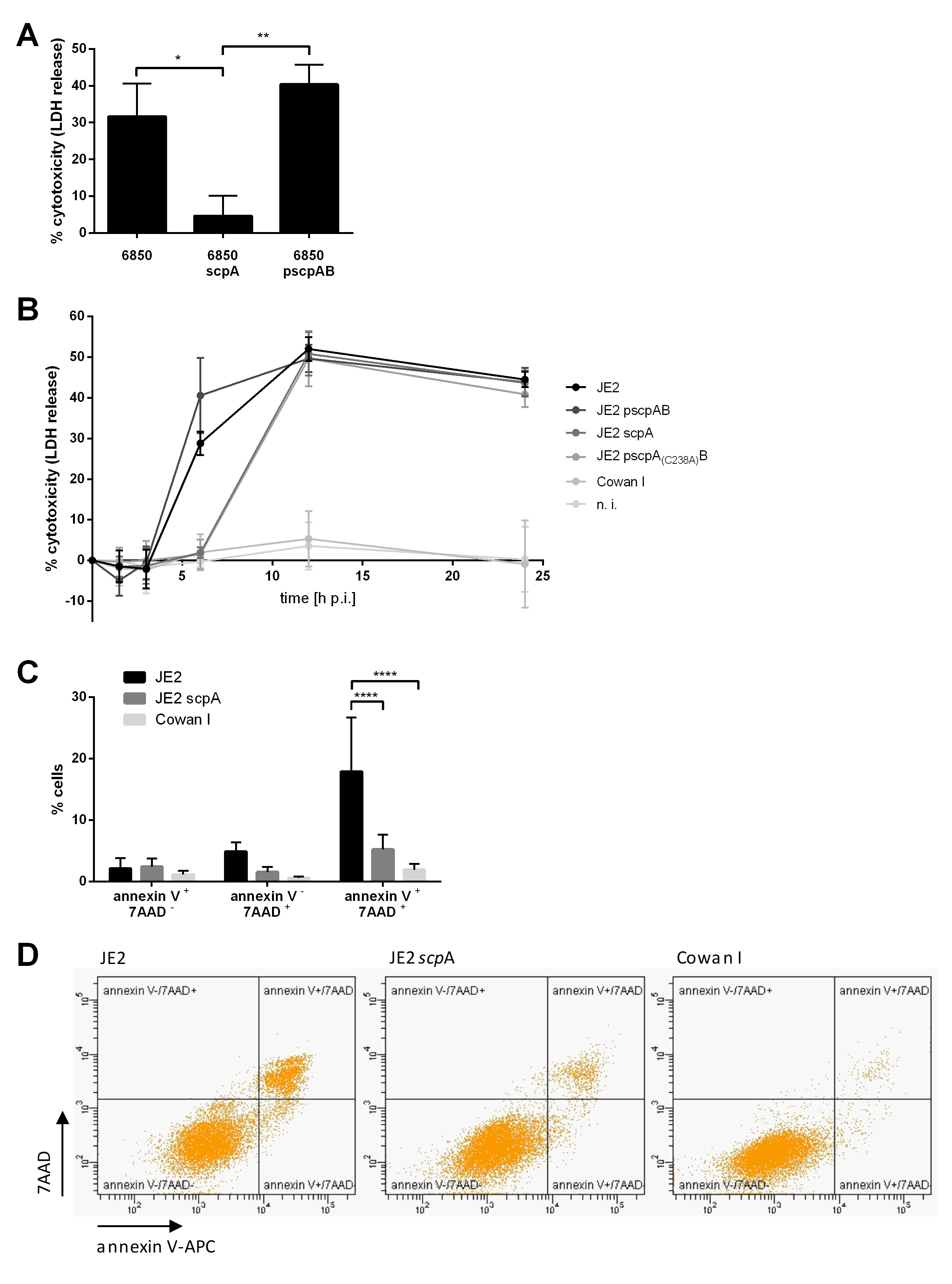

### Supplemental Figure 2

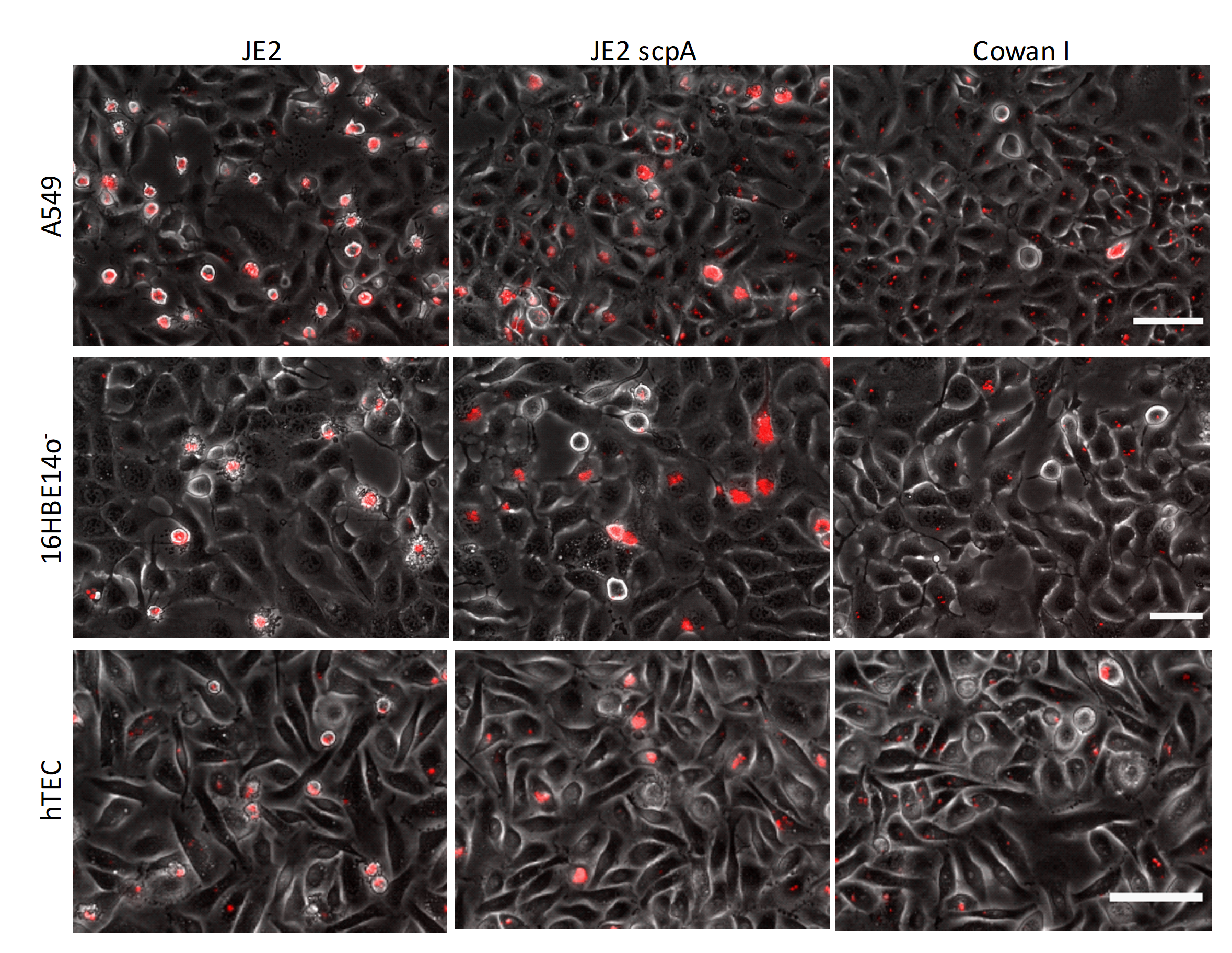

### Supplemental Figure 3

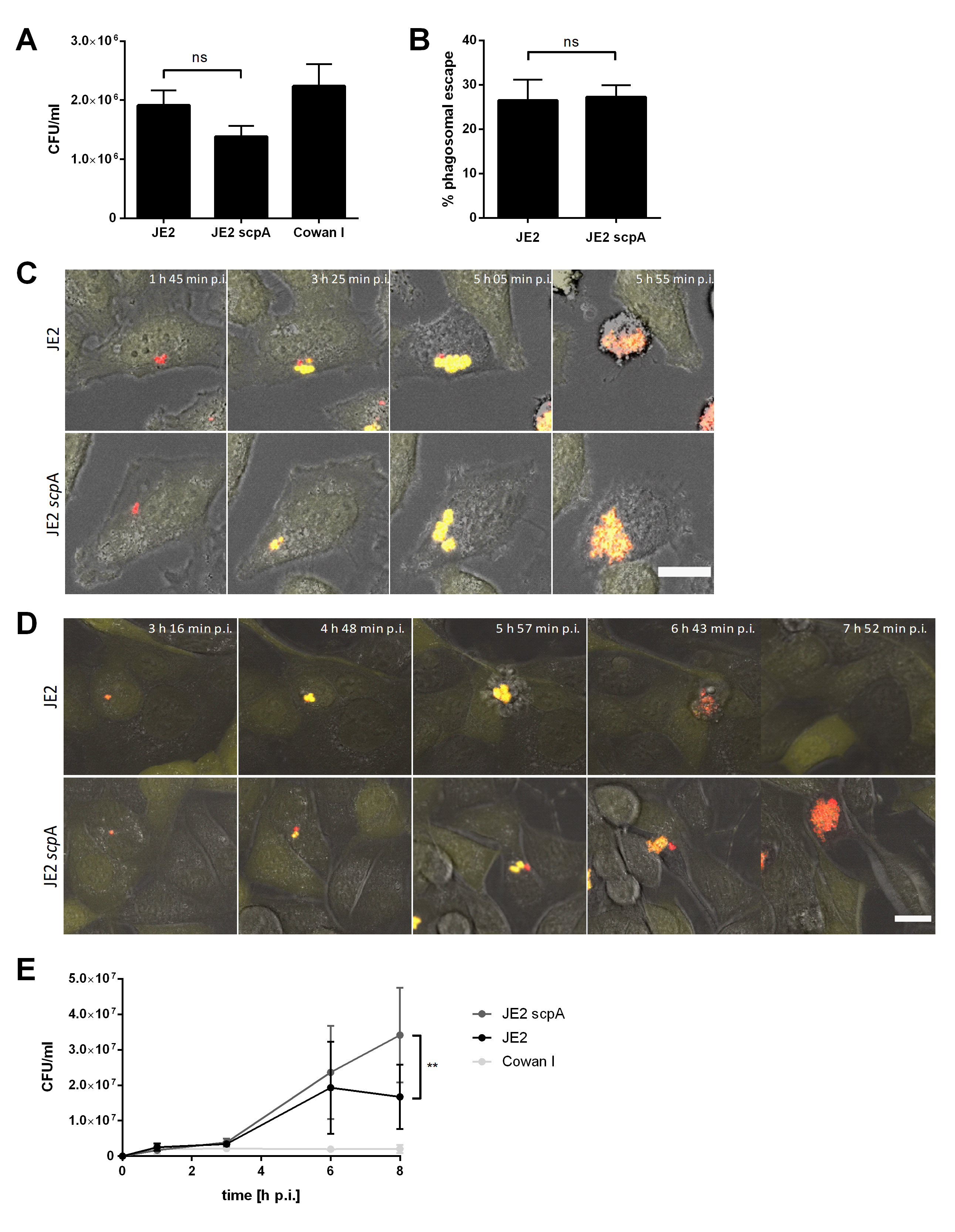

### Supplemental Figure 4

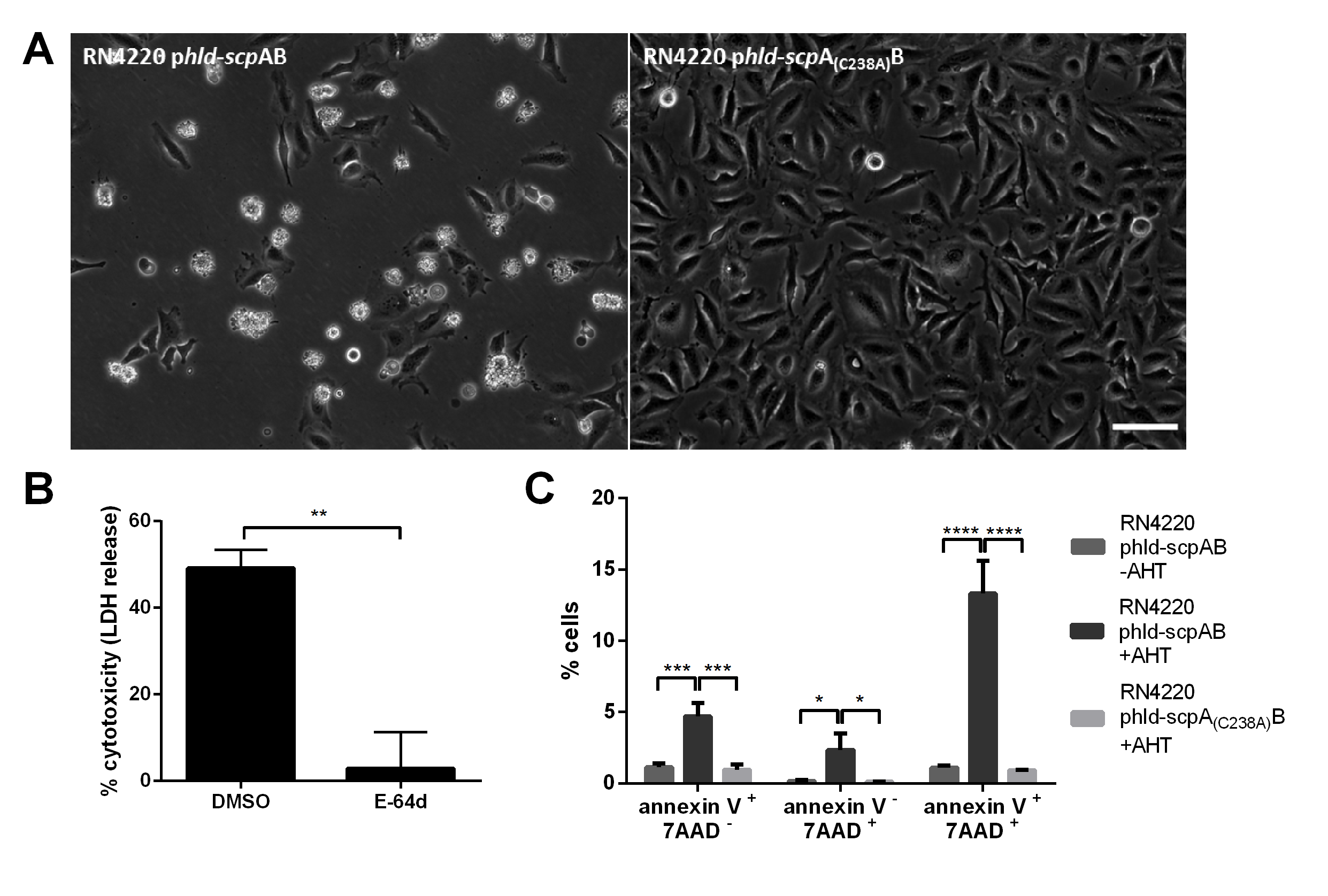

### Supplemental Figure 5

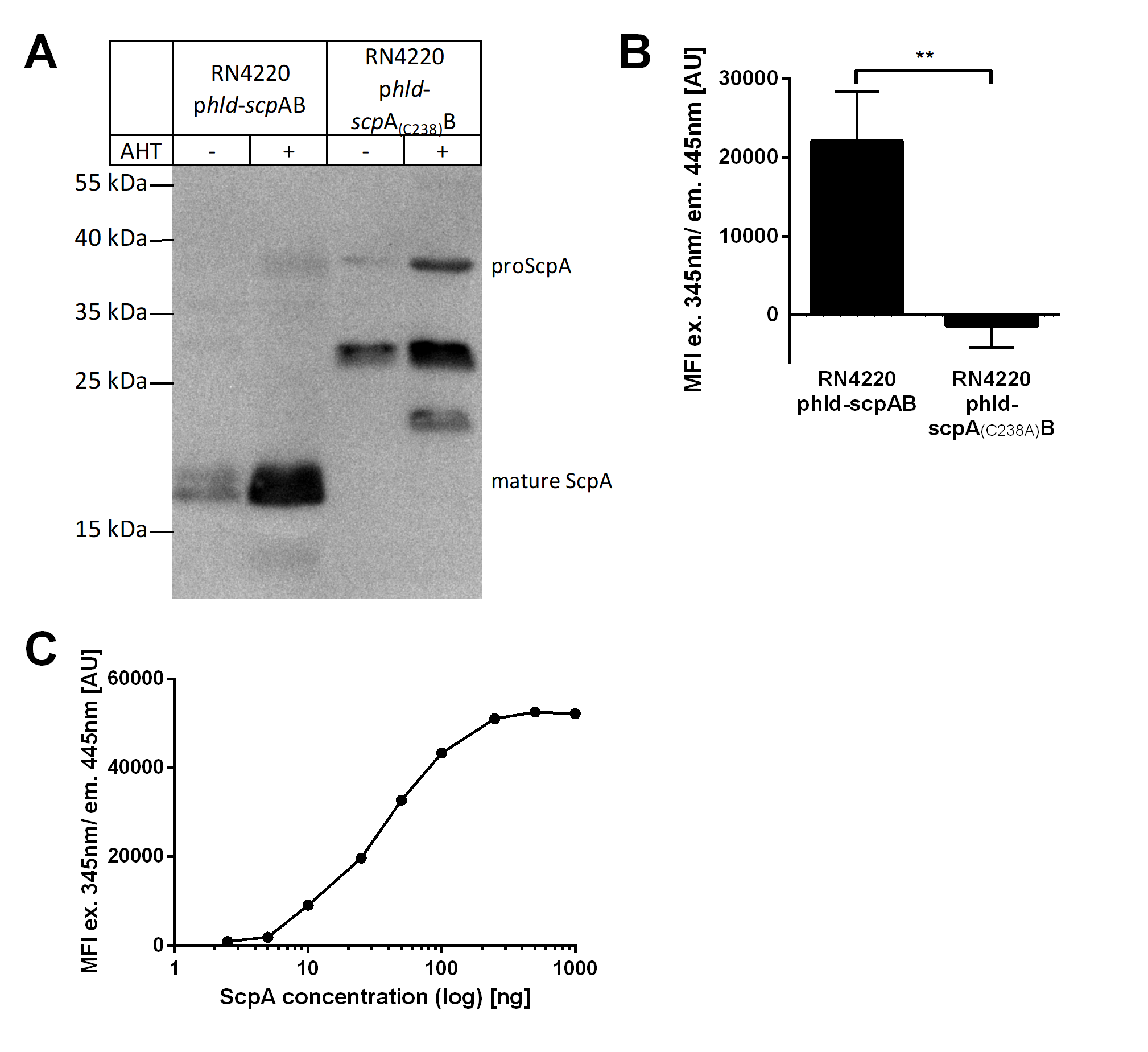

### Supplemental Figure 6

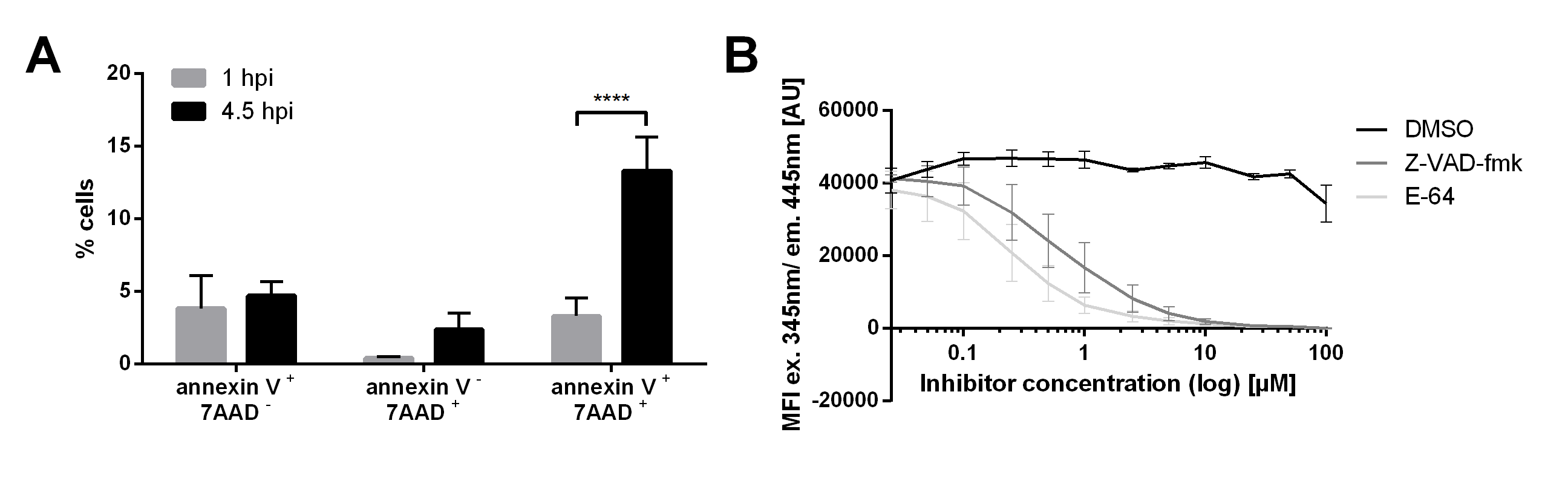

### Supplemental Figure 7

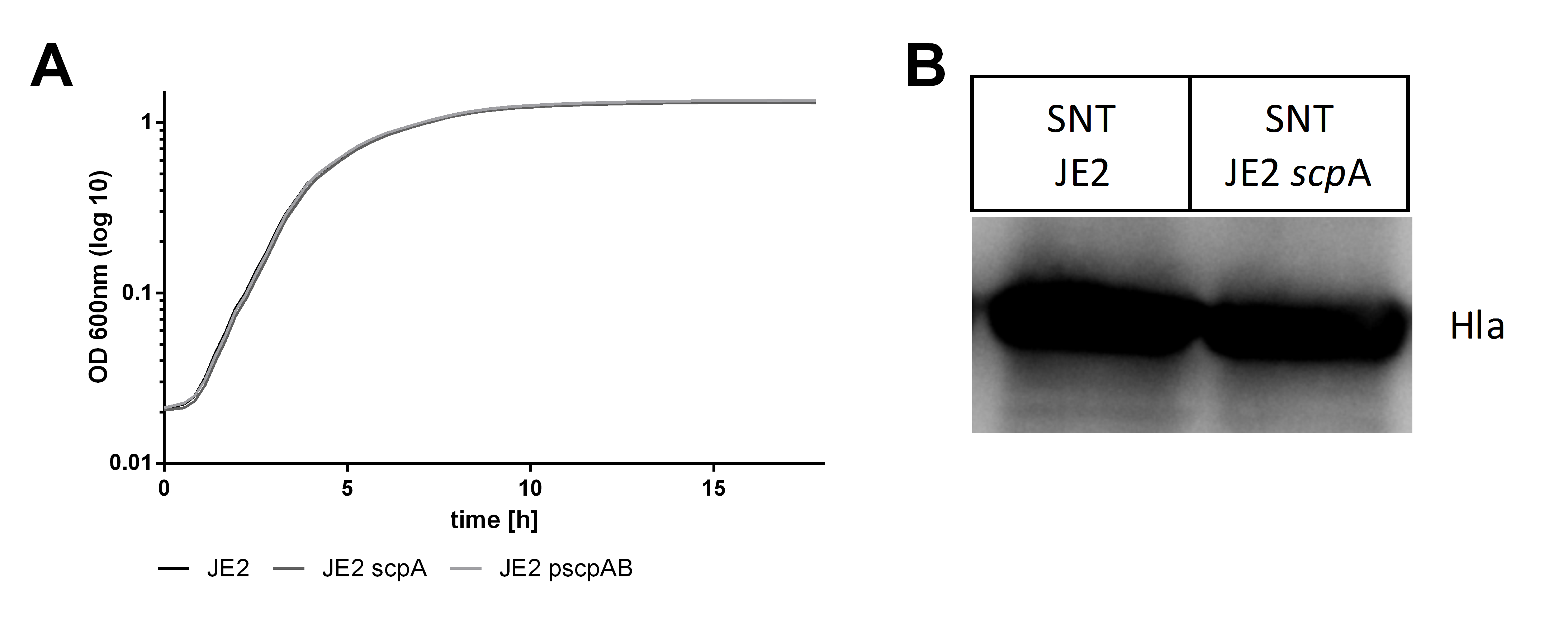
